# Covalent tumor anchoring spatially orchestrates antitumor immunity

**DOI:** 10.64898/2026.05.13.724746

**Authors:** Qingke Li, Hongfei Chen, Pan Zhang, Li Cao, Bingchen Yu, Lei Wang

## Abstract

Protein immunotherapies can elicit potent tumor rejection, but reversible target engagement, incomplete tumor retention, and systemic leakage often erode spatial control. Here, we develop covalently anchored tumor immunotherapeutic proteins (CATIPs), a modular platform that uses proximity-enabled covalent chemistry to immobilize immune cues on tumor-cell surfaces after intratumoral administration. CATIPs combine tumor-targeting nanobodies with payloads for T cell engagement, co-stimulation, and cytokine support. In human PBMC-reconstituted NSG mice, CATIPs completely eradicated treated EGFR-positive tumors, outperforming matched non-covalent proteins while limiting redistribution, systemic T cell activation, cytokine release, xGVHD-associated morbidity, and on-target, off tumor toxicity. In immunocompetent melanoma models, CATIPs remodeled the tumor microenvironment, expanded antigen-specific CD8^+^ T cells, induced antigen-restricted abscopal control, and generated durable protection against local and metastatic rechallenge. CATIP-engineered tumor cells further functioned as whole-cell vaccines. Thus, covalent tumor anchoring converts local protein delivery into tumor-surface immune programming, enabling systemic, tumor-specific, durable antitumor immunity while limiting systemic immunopathology.

## INTRODUCTION

A central challenge in cancer immunotherapy is to intensify immune activation within tumors without distributing that activity systemically.^1,2^ Physiological T cell immunity is protected from indiscriminate activation by anatomical and cellular compartmentalization: naive T cells are primed in secondary lymphoid organs by antigen-presenting cells that coordinate antigen recognition with co-stimulation and cytokine cues, whereas effector T cells are restimulated in inflamed tissues through local cell–cell encounters.^3,4^ Many cancer immunotherapies uncouple T cell activation from this spatial control. T cell engagers, agonistic antibodies, cytokines, immune checkpoint inhibitors, and adoptive cell therapies can drive tumor rejection, but when their activity extends beyond tumor tissue, the same signals can provoke cytokine release, immune-related adverse events and on-target, off-tumor (OTOT) injury in normal tissues expressing the targeted antigen.^5–11^ This trade-off is especially limiting in solid tumors, where poor immune infiltration, antigen heterogeneity and immunosuppressive tumor microenvironments demand potent local immune activation, yet systemic exposure narrows the therapeutic window.^3,4,12,13^ Intratumoral administration can enrich immunotherapy at the tumor site and is clinically feasible for many tumors, but most locally delivered proteins remain reversible and diffusible.^13,14^ As a result, active molecules can dissociate from tumor cells, redistribute beyond the injected tumor and lose the spatial coupling required for sustained immune signaling.^13,15–17^

This limitation is particularly important for protein immunotherapies whose activity depends on spatially organized cell-cell signaling. Soluble T cell engagers must form transient complexes among a tumor antigen, CD3 and the engager itself, a process constrained by dissociation, diffusion and non-productive receptor occupancy at high exposure.^7,18^ Cytokines and immune agonists similarly require sufficient local persistence to remodel the tumor microenvironment, yet systemic exposure can rapidly convert immune stimulation into toxicity.^9,15–17,19,20^ Existing localization approaches, including collagen anchoring, biomaterial depots, conditional activation, cell-surface targeting, and intratumoral expression via engineered T cells, demonstrate that spatial confinement can improve the therapeutic index.^15–17,21–25^ However, these approaches generally rely on non-covalent interactions and therefore do not durably couple immunostimulatory signals to tumor cells. Whether potent combinatorial immune inputs can be stably assembled at tumor-cell surfaces, preserving local organization while initiating systemic antitumor immunity, remains unclear.

We reasoned that covalent tumor anchoring could convert local delivery into durable tumor-surface immune programming. Proximity-enabled reactive therapeutics (PERx) use genetically encoded latent bioreactive amino acids that form covalent bonds only after binding, enabling selective and irreversible capture of native protein targets.^26–28^ Applied to tumor-targeted immunomodulators, this covalent chemistry should allow immunostimulatory proteins to remain fixed on tumor cells, sustain signaling at sites of immune-cell contact and limit systemic escape. In this way, covalent anchoring could functionally recouple key elements of T cell activation by enforcing coordinated receptor engagement at spatially restricted cellular interfaces. It also provides a route to impose antigen-presenting-cell-like functions directly onto tumor cells by displaying defined combinations of T cell engagement, co-stimulation, cytokine support and checkpoint-modulating modules on the tumor membrane, without genetic manipulation of either tumor cells or lymphocytes.^17,29–31^

Here we develop covalently anchored tumor immunotherapeutic proteins (CATIPs) as a modular platform for spatially restricted local immunotherapy. CATIPs combine a tumor-targeting nanobody containing a latent bioreactive amino acid with immunomodulatory payloads that deliver key signals for immune activation. We show that intratumorally administered CATIPs remain associated with tumor cells, outperform matched non-covalent proteins, eradicate established tumors, and limit systemic T cell activation and off-tumor toxicity. In immunocompetent melanoma models, CATIPs induce antigen-specific systemic immunity and durable protective memory, and enable tumor cells to function as whole-cell vaccines that confer tumor-specific protection. These findings define covalent tumor anchoring as a general strategy to spatially couple potent immune signals to tumor cells, providing a framework for local immunotherapy that elicits systemic antitumor immunity while limiting systemic immunopathology.

## RESULTS

### Design and engineering of CATIPs

To achieve stable localization of immunostimulatory proteins to tumor cells, we designed CATIPs as modular fusion proteins in which a tumor-targeting nanobody is linked to an immunomodulatory payload through a flexible (GGGGS)_4_ linker (**Fig. 1A**). The targeting module recognizes receptors overexpressed on tumor cells, whereas the effector modules provide the three canonical inputs required for T cell activation: CD3 engagement for TCR triggering (signal 1), CD28 engagement for co-stimulation (signal 2), and cytokine receptor engagement (signal 3). To enable irreversible tumor binding, we incorporated the latent bioreactive Uaa FSY into the nanobody,^32^ allowing site-specific covalent receptor capture through PERx mechanism.^26^ This modular design permits the use of individual CATIPs or defined CATIP combinations.

**Figure 1.**
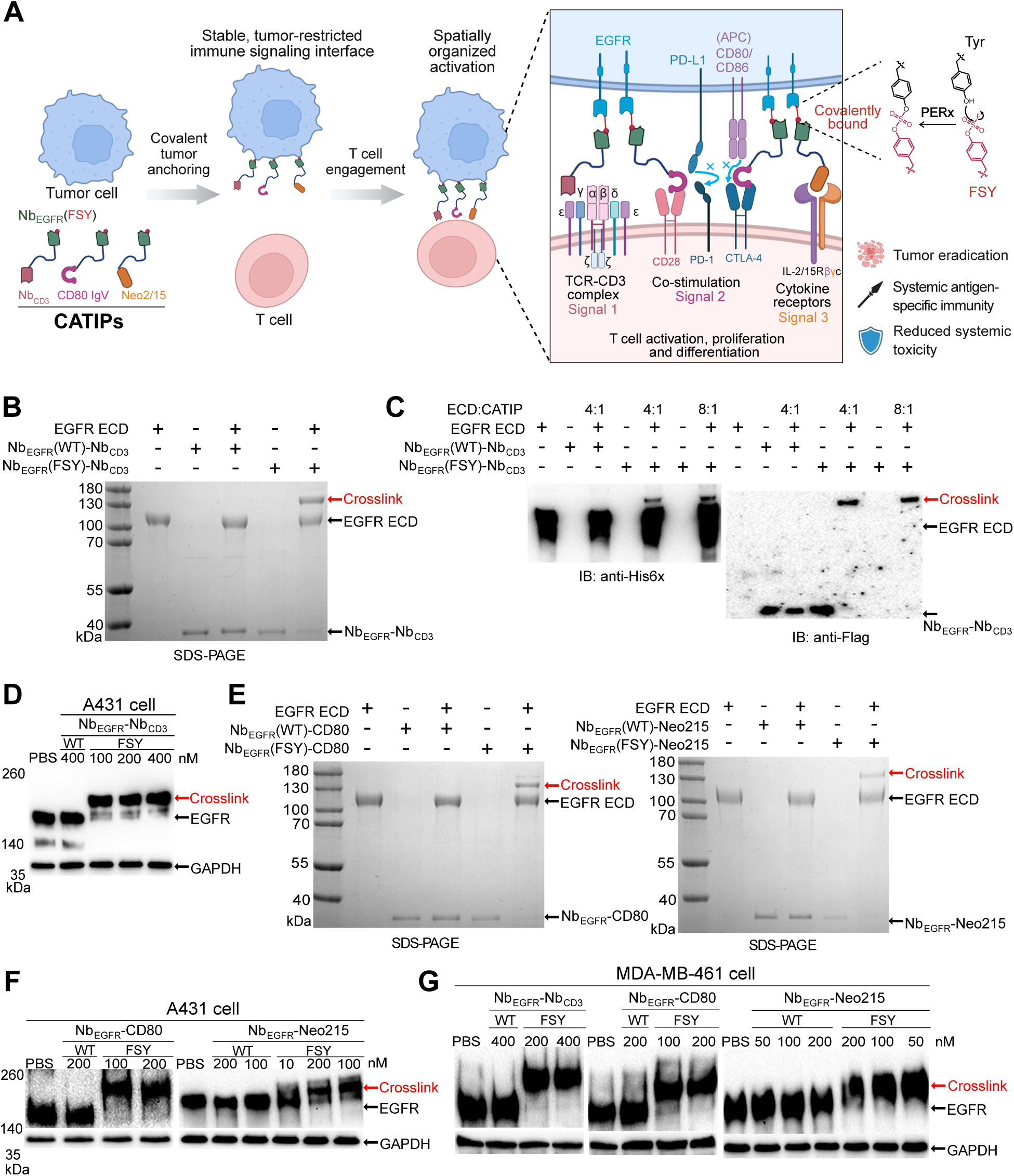
CATIPs covalently crosslink EGFR in vitro and on tumor cell surfaces. (A) Concept of covalent tumor anchoring by CATIPs. Conventional immunotherapies rely on reversible interactions and can dissociate and redistribute beyond the tumor microenvironment. CATIPs use proximity-enabled covalent chemistry to immobilize immunostimulatory proteins on tumor-cell surfaces after binding, enabling stable and spatially confined presentation of T cell engagement, co-stimulation, and cytokine support. This covalent anchoring creates a persistent, membrane-confined immune signaling interface that promotes localized T cell activation and enables antigen-specific systemic immunity with reduced systemic exposure and toxicity. (B) Nb_EGFR_(FSY)-Nb_CD3_ covalently crosslinked EGFR extracellular domain (ECD) in vitro. Nb_EGFR_-Nb_CD3_ was incubated with EGFR ECD at 1:1 molar ratio in PBS at 37°C for 16 h. (C) Increasing the EGFR ECD:Nb_EGFR_(FSY)–Nb_CD3_ molar ratio to 4:1 or 8:1 resulted in near-complete covalent crosslinking of Nb_EGFR_(FSY)–Nb_CD3_ in PBS at 37°C for 12 h. (D) Nb_EGFR_(FSY)–Nb_CD3_ covalently crosslinked full-length EGFR on the surface of A431 cells. (E) Nb_EGFR_(FSY)-CD80 (left) and Nb_EGFR_(FSY)-Neo2/15 (right) efficiently crosslinked EGFR ECD in vitro at a 1:1 molar ratio in PBS at 37°C for 16 h. (F) Nb_EGFR_(FSY)-CD80 and Nb_EGFR_(FSY)-Neo2/15 covalently crosslinked EGFR on the surface of A431 cells. (G) Nb_EGFR_(FSY)–Nb_CD3_, Nb_EGFR_(FSY)-CD80, and Nb_EGFR_(FSY)-Neo2/15 covalently crosslinked EGFR on the surface of MDA-MB-468 cells.

Given the frequent overexpression of ErbB family receptors in solid tumors and their established roles in tumorigenesis and immune evasion,^33^ we first generated FSY-containing nanobodies targeting HER-family receptors. Introduction of Arg at selected positions further accelerated covalent crosslinking.^34^ Covalent targeting of HER2 led to marked receptor degradation on tumor cells (**Fig. S1**), consistent with the known rapid internalization of HER2 after antibody engagement.^35^ In contrast, EGFR crosslinking did not produce significant degradation (see below). We therefore focused on EGFR and generated an EGFR-targeted covalent nanobody, Nb_EGFR_(FSY), which mediated rapid, time-dependent crosslinking of EGFR.^34^

To deliver signal 1, we fused Nb_EGFR_(FSY) to a human CD3ε agonistic nanobody,^36^ generating Nb_EGFR_(FSY)–Nb_CD3_. This construct efficiently crosslinked the EGFR extracellular domain (ECD) in vitro (**Fig. 1B**). At EGFR ECD:Nb_EGFR_(FSY)-Nb_CD3_ molar ratios of 4:1 and 8:1, crosslinking was nearly complete (**Fig. 1C**). In cells, 100 nM Nb_EGFR_(FSY)-Nb_CD3_ crosslinked endogenous full-length EGFR on A431 cells almost quantitatively, whereas the wild-type control, Nb_EGFR_(WT)-Nb_CD3_, produced no detectable crosslinking (**Fig. 1D**).

To provide signal 2, we generated Nb_EGFR_(FSY)–CD80 by fusing Nb_EGFR_(FSY) to a CD80 IgV variant that mediates PD-L1 binding–dependent CD28 co-stimulation.^37,38^ As reported, this variant binds PD-L1 and CTLA-4, antagonizing PD-1–PD-L1 and CTLA-4–CD80/CD86 inhibitory pathways and potentially increasing the availability of endogenous CD80/CD86 on APCs, thereby enhancing CD28-mediated co-stimulation.^38^ To deliver signal 3, we generated Nb_EGFR_(FSY)–Neo2/15, incorporating a de novo designed IL-2/IL-15 mimic that binds the IL-2 receptor βγ_c_ heterodimer (IL-2Rβγ_c_) but not IL-2Rα or IL-15Rα, minimizing preferential stimulation of IL-2Rα–high regulatory T cells.^39^

Both Nb_EGFR_(FSY)–CD80 and Nb_EGFR_(FSY)–Neo2/15 efficiently crosslinked the EGFR ECD in vitro (**Fig. 1E)** and robustly crosslinked endogenous EGFR on A431 cells (**Fig. 1F**). All three CATIPs also efficiently crosslinked endogenous EGFR on MDA-MB-468 cells (**Fig. 1G**). These data establish a modular CATIP platform capable of delivering the three principal inputs required for T cell activation while achieving efficient covalent anchoring to EGFR in vitro and on the tumor cell surface.

### CATIP retention and distribution in cells and in vivo

We next examined whether CATIPs remain associated with cells after covalently anchoring. Tumor cells were incubated CATIPs, and culture supernatants were collected at the indicated time points for western blot analysis. Nb_EGFR_(WT)–Nb_CD3_, which binds EGFR non-covalently, was detected predominantly in the supernatant and remained largely unchanged over time, consistent with rapid dissociation from the cell surface. In contrast, the amount of Nb_EGFR_(FSY)–Nb_CD3_ in the supernatant progressively decreased under both conditions tested: a fixed cell number with increasing protein concentrations (10 nM and 50 nM) (**Fig. 2A**), and a fixed protein concentration (25 nM) with increasing cell numbers (0.5, 2.5, or 10 × 10⁶ cells) (**Fig. 2B**). These data indicate progressive anchoring of Nb_EGFR_(FSY)–Nb_CD3_ on tumor cells.

**Figure 2.**
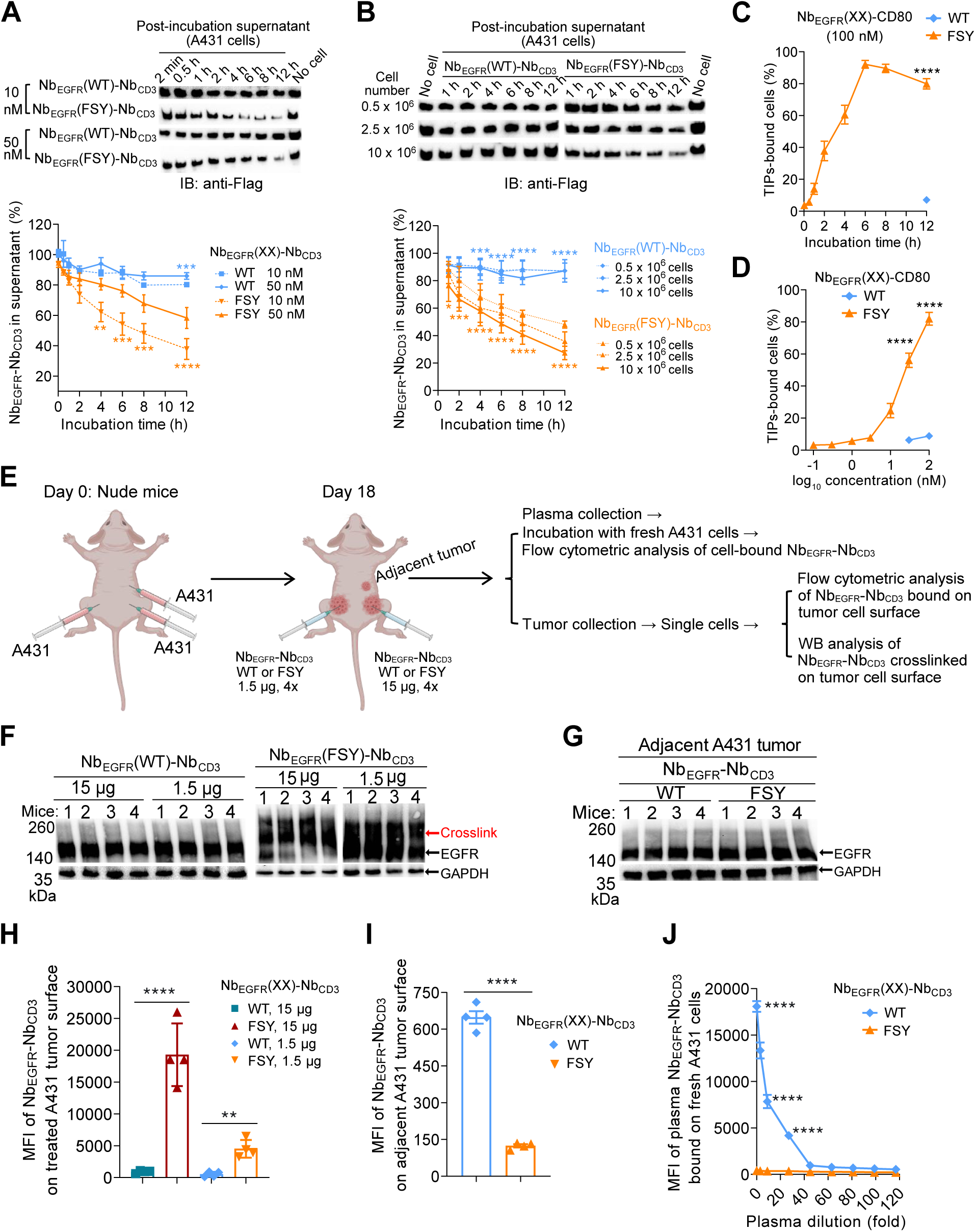
CATIPs are retained on tumor cells and remained localized in vivo. (A,B) Nb_EGFR_(FSY)–Nb_CD3_ was progressively depleted from culture supernatants after incubation with A431 cells, either at fixed number with increasing protein concentrations (**A**) or at fixed protein concentration with increasing cell number (**B**), consistent with covalent anchoring. Data are mean ± SEM. (**A**) n = 7 independent experiments; (**B**) n = 3 independent experiments for Nb_EGFR_(WT)-Nb_CD3_ at 2.5 x 10^6^ cells, n = 5 at 0.5 x and 10 x 10^6^ cells, and n = 6 for Nb_EGFR_(FSY)-Nb_CD3_. \**P* < 0.05, ** *P* < 0.01, *** *P* < 0.001, **** *P* < 0.0001; asterisks at the indicated positions denote comparisons between WT and FSY proteins at matched concentrations or cell numbers (two-way ANOVA with Tukey’s multiple-comparisons test). **(C,D)** Nb_EGFR_(FSY)–CD80 showed time-dependent (**C**) and concentration-dependent (**D**) retention on A431 cells after PBS washing, consistent with stable cell-surface anchoring. Data are mean ± SEM. n = 4 (**C**) or 5 (**D**) independent experiments. **** *P* < 0.0001; unpaired *t*-test (C) or two-way ANOVA with Šídák’s multiple-comparisons test (D). (E) Experimental scheme for in vivo evaluation of CATIP retention and distribution. (F) Intratumorally administered Nb_EGFR_(FSY)–Nb_CD3_ covalently crosslinked EGFR in treated A431 tumors in a dose-dependent manner. (G) No detectable EGFR crosslinking was observed in untreated adjacent A431 tumors. **(H,I)** Flow cytometric analysis of the mean fluorescence intensity (MFI) of Nb_EGFR_(FSY)–Nb_CD3_ on tumor cells from treated (**H**) or untreated adjacent (**I**) A431 tumors. Data are mean ± SEM. n = 4. \*\**P* < 0.01; **** *P* < 0.0001; unpaired *t*-test. For (H), low-dose and high-dose TIPs groups were analyzed independently and then combined for presentation. (J) Flow cytometric analysis of the MFI of plasma-derived Nb_EGFR_(FSY)–Nb_CD3_ bound to fresh A431 cells in vitro. Data are mean ± SEM. n = 4. \*\*\*\**P* < 0.0001; two-way ANOVA with Šídák’s multiple-comparisons test.

Similar results were obtained with Nb_EGFR_–CD80. After incubation with tumor cells followed by PBS washing, cell-associated protein was quantified by flow cytometry. Whereas Nb_EGFR_(WT)–CD80 showed residual binding after washing, the fraction of cells positive for Nb_EGFR_(FSY)–CD80 increased with both incubation time and protein concentration, consistent with stable anchoring on tumor cells (**Fig. 2C, D**). Similar behavior was observed for Nb_EGFR_(FSY)–Neo2/15 (**Fig. S2**). Together, these in vitro data indicate that CATIPs bind EGFR irreversibly and remain cell associated after washing, whereas the corresponding WT proteins dissociate and redistribute.

We next asked whether intratumorally administered Nb_EGFR_(FSY)–Nb_CD3_ could crosslink human EGFR in tumor tissue in situ and whether it could redistribute to crosslink EGFR outside the treated tumor. NSG mice bearing bilateral A431 tumors received intratumoral injections of Nb_EGFR_(FSY)-Nb_CD3_ into one tumor at either a low dose (1.5 μg) or a high dose (15 μg) every 12 hours for a total of 4 doses. A separate adjacent A431 tumor was left untreated as a control for off-site EGFR crosslinking (**Fig. 2E)**. Tumors were harvested and analyzed for EGFR crosslinking by western blot. In treated tumors, Nb_EGFR_(WT)–Nb_CD3_ produced no detectable EGFR crosslinking at either dose (**Fig. 2F**). In contrast, Nb_EGFR_(FSY)–Nb_CD3_ induced robust EGFR crosslinking at both doses, with greater crosslinking at the higher dose (**Fig. 2F**).

No EGFR crosslinking was detected in the untreated adjacent A431 tumors, regardless of whether mice received Nb_EGFR_(WT)–Nb_CD3_ or Nb_EGFR_(FSY)–Nb_CD3_, indicating that intratumorally administered Nb_EGFR_(FSY)–Nb_CD3_ did not measurably diffuse to adjacent tumors to mediate off-site covalent crosslinking (**Fig. 2G**). Consistent with this finding, flow cytometric analysis of tumor cell-associated Nb_EGFR_–Nb_CD3_ showed markedly greater retention of Nb_EGFR_(FSY)–Nb_CD3_ than Nb_EGFR_(WT)–Nb_CD3_ in treated tumors, reaching 21.2-fold at the high dose and 8.7-fold at the low dose (**Fig. 2H**). In contrast, binding of Nb_EGFR_(WT)–Nb_CD3_ in untreated A431 tumors was 5.5-fold higher than that of Nb_EGFR_(FSY)–Nb_CD3_ (**Fig. 2I**). To assess systemic redistribution, we measured Nb_EGFR_–Nb_CD3_ in plasma by incubating collected plasma with fresh A431 cells, followed by flow cytometric detection of cell-bound Nb_EGFR_–Nb_CD3_ (**Fig. 2J**). Substantial amounts of Nb_EGFR_(WT)–Nb_CD3_ were detected in plasma, at 46-fold above Nb_EGFR_(FSY)–Nb_CD3_, whereas Nb_EGFR_(FSY)–Nb_CD3_ remained at baseline.

Overall, these in vivo data indicate that intratumorally administered Nb_EGFR_(WT)–Nb_CD3_ diffused into the circulation and redistributed, whereas Nb_EGFR_(FSY)–Nb_CD3_ was retained predominantly within the treated tumor through covalent anchoring, thereby limiting off-site EGFR crosslinking in untreated tumors.

### CATIPs enhance T cell activation in vitro

To determine whether covalent anchoring of Nb_EGFR_(FSY)–Nb_CD3_ enhances T cell activation, we performed in vitro PBMC–tumor cell co-culture assays. Pre-seeded A431 cells were treated with either Nb_EGFR_(WT)–Nb_CD3_ or Nb_EGFR_(FSY)–Nb_CD3_ for 12 h, followed by either PBS washing or no washing before co-cultured with PBMCs. Under the PBS-washed condition, Nb_EGFR_(FSY)–Nb_CD3_ markedly enhanced PBMC-mediated killing of A431 cells after 72 h, with substantial activity already evident at 5 nM and a concentration-dependent increase in cytotoxicity (**Fig. 3A**). By contrast, Nb_EGFR_(WT)–Nb_CD3_ induced only ∼25% killing even at 100 nM. Under the non-washed condition, Nb_EGFR_(FSY)–Nb_CD3_ again elicited substantially greater PBMC-mediated cytotoxicity than Nb_EGFR_(WT)–Nb_CD3_ after 48 h, with an approximately 10-fold improvement in EC_50_ (**Fig. 3B**).

**Figure 3.**
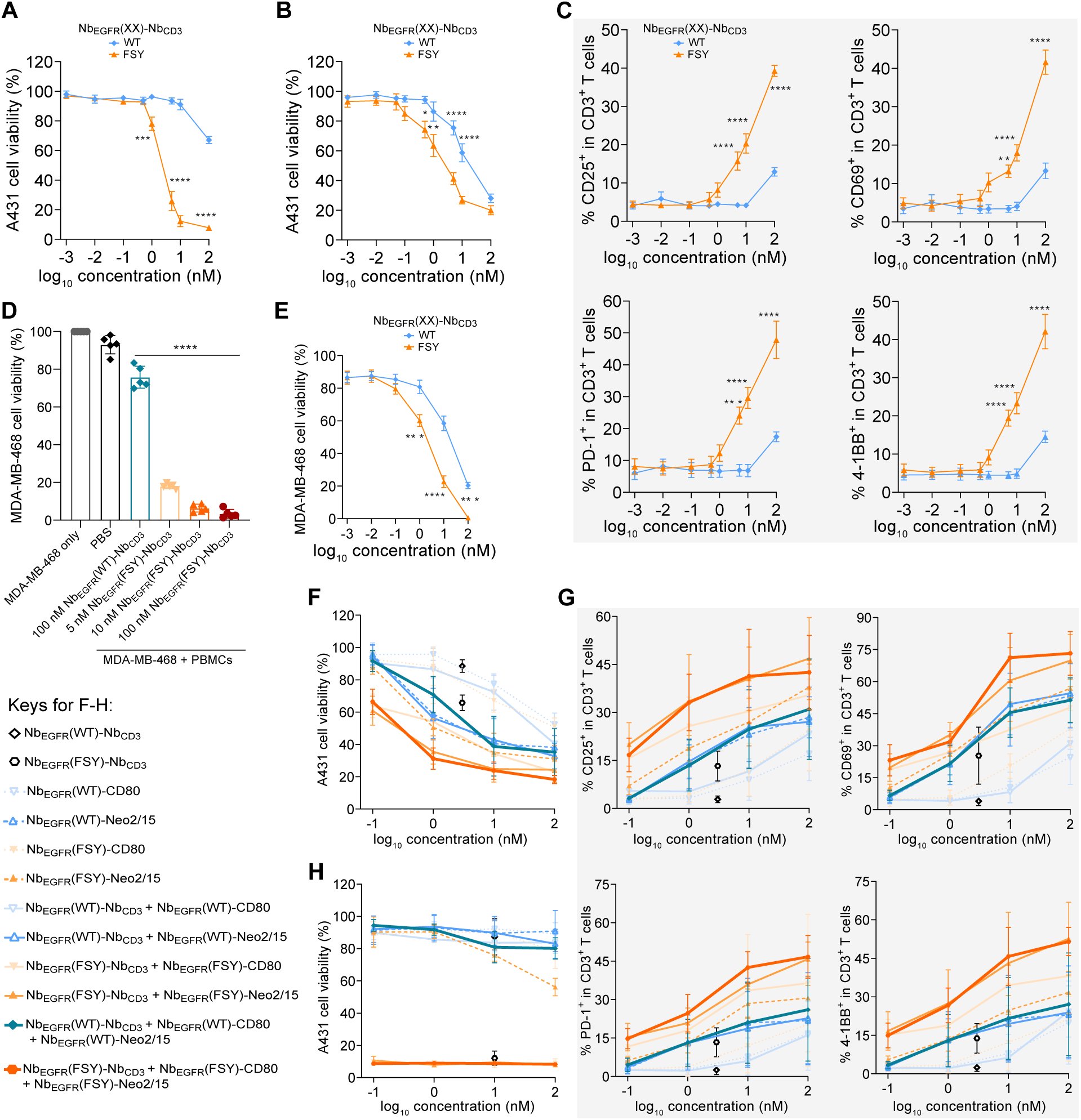
CATIPs enhance T cell activation and PBMC-mediated tumor cell cytotoxicity. (A-B) Nb_EGFR_(FSY)–Nb_CD3_ enhanced PBMC-mediated killing of A431 cells under PBS-washed (A) or non-washed (**B**) conditions in a dose-dependent manner. Data are mean ± SEM. n = 4 independent experiments. \**P* < 0.05; \*\**P* < 0.01; \*\*\**P* < 0.001; \*\*\*\**P* < 0.0001; two-way ANOVA with Šídák’s multiple-comparisons test. **(C)** Nb_EGFR_(FSY)–Nb_CD3_ increased expression of the indicated T cell activation markers in a dose-dependent manner. A431 cells were pre-incubated with Nb_EGFR_(FSY)–Nb_CD3_ under PBS-washed conditions and then co-cultured with PMBCs. Data are mean ± SEM. n = 4 independent experiments. \*\**P* < 0.01; \*\*\**P* < 0.001; \*\*\*\**P* < 0.0001; two-way ANOVA with Šídák’s multiple-comparisons test. **(D-E)** Nb_EGFR_(FSY)–Nb_CD3_ enhanced PBMC-mediated cytotoxicity against MDA-MB-468 cells under PBS-washed (**D**) or non-washed (**E**) conditions in a dose-dependent manner. Data are mean ± SEM. n = 5 independent experiments. \*\*\**P* < 0.001, \*\*\*\**P* < 0.0001; one-way ANOVA with Tukey’s multiple-comparisons test (**D**) or two-way ANOVA with Šídák’s multiple-comparisons test (**E**). **(F)** Under non-washed conditions, CATIP combinations cooperatively enhanced PBMC-mediated cytotoxicity against A431 cells in a dose-dependent manner. The final concentration of Nb_EGFR_–Nb_CD3_ was 3 nM. **(G)** Under non-washed conditions, CATIP combinations cooperatively increased expression of T cell activation markers CD25, CD69, PD-1 and 4-1BB. A431 cells were pre-incubated with CATIPs and then co-cultured with PBMCs. The final concentration of Nb_EGFR_–Nb_CD3_ was 3 nM. **(H)** Under PBS-washed conditions, CATIP combinations induced robust cooperative PBMC-mediated cytotoxicity against A431 cells, whereas corresponding WT combinations were largely ineffective. The final concentration of Nb_EGFR_–Nb_CD3_ was 10 nM.

We next assessed T cell activation by flow cytometric analysis of activation markers. Consistent with the cytotoxicity data, Nb_EGFR_(FSY)–Nb_CD3_ at 5 nM significantly increased expression of CD25, CD69, PD-1 and 4-1BB under the PBS-washed condition, with further increase at higher concentrations (**Fig. 3C**). By contrast, Nb_EGFR_(WT)–Nb_CD3_ at 100 nM induced only modest upregulation of these markers. Similar trends were observed under the non-washed condition (**Fig. S3A)**.

Nb_EGFR_(FSY)–Nb_CD3_ likewise enhanced PBMC-mediated cytotoxicity against MDA-MB-468 cells under both PBS-washed (**Fig. 3D**) and non-washed conditions (**Fig. 3E**), and similarly increased T cell activation in both settings (**Fig. S3B, C)**. These results show that covalent anchoring to EGFR substantially enhances the activity of Nb_EGFR_(FSY)–Nb_CD3_, resulting in stronger CD3ε-mediated T cell activation and greater cytotoxicity than the non-covalent construct across both tumor cell lines.

We next examined the cooperative effects of Nb_EGFR_(FSY)–Nb_CD3_ with Nb_EGFR_(FSY)–CD80 and Nb_EGFR_(FSY)–Neo2/15 on PBMC–mediated tumor cell killing and T cell activation. Because CD28 co-stimulation by the CD80 IgV variant depends on its interaction with PD-L1, we first confirmed that both A431 and MDA-MB-468 cells upregulate PD-L1 in response to IFNγ stimulation (**Fig. S3D**). Pre-seeded A431 cells were then incubated with Nb_EGFR_(FSY)–Nb_CD3_, Nb_EGFR_(FSY)–CD80 and/or Nb_EGFR_(FSY)–Neo2/15 for 12 h before addition of PBMCs.

At 15 nM Nb_EGFR_–Nb_CD3_, Nb_EGFR_(FSY)–Nb_CD3_ alone induced ∼70% tumor cell killing (**Fig. S3E**). Under these conditions, addition of Nb_EGFR_(FSY)–CD80 or Nb_EGFR_(FSY)–Neo2/15 yielded only limited further increases in killing, indicating that the activity of Nb_EGFR_(FSY)–Nb_CD3_ alone was already near maximal (**Fig. S3E**). By contrast, the corresponding combinations of Nb_EGFR_(WT)–Nb_CD3_ with Nb_EGFR_(WT)–CD80 and/or Nb_EGFR_(WT)–Neo2/15 increased tumor cell killing from ∼25% to ∼55%. Consistent with these, T-cell activation markers in the Nb_EGFR_(FSY)–Nb_CD3_ alone group were already elevated to ∼80%, leaving little headroom for further enhancement, whereas the WT combinations showed clear cooperative increases in activation (**Fig. S3F**). Even so, both cytotoxicity and T cell activation remained greater in the CATIP groups than in the corresponding WT-TIP groups.

At a lower concentration of Nb_EGFR_–Nb_CD3_ (3 nM), Nb_EGFR_(FSY)–Nb_CD3_ alone induced ∼35% killing (**Fig. 3F**). Under these conditions, combining Nb_EGFR_(FSY)–Nb_CD3_ with Nb_EGFR_(FSY)–CD80 and/or Nb_EGFR_(FSY)–Neo2/15 markedly enhanced cytotoxicity, increasing killing to ∼80% (**Fig. 3F**). By contrast, the corresponding WT combinations showed substantially lower activity and did not outperform individual components or the two-component combinations. Consistent with the cytotoxicity data, FSY-containing combinations markedly increased the expression of CD25, CD69, PD-1 and 4-1BB (**Fig. 3G**), whereas WT three-component combinations did not significantly enhance T cell activation relative to single agents or two-component combinations.

Notably, under PBS-washed conditions, all two- and three-component CATIP combinations induced >90% tumor cell killing, whereas the corresponding WT combinations left ∼85% of tumor cells viable (**Fig. 3H**).

These data indicate that covalent EGFR targeting by Nb_EGFR_(FSY)–Nb_CD3_ enhances PBMC-mediated cytotoxicity and T cell activation and supports cooperative effects with Nb_EGFR_(FSY)–CD80 and Nb_EGFR_(FSY)–Neo2/15, particularly under PBS-washed conditions in which the non-covalent constructs are largely ineffective.

### CATIPs induce complete tumor clearance with minimal toxicity

We next examined whether covalent anchoring improves antitumor efficacy and reduces OTOT toxicity in vivo (**Fig. 4A**). To approximate human anti-tumor immune responses, we used immunodeficient NSG mice adoptively transferred with human PBMCs (hPBMCs). This hPBMC-humanized NSG model enables concurrent assessment of antitumor efficacy and therapy-associated immune toxicities.^40–43^ We established a local therapeutic tumor using the human EGFR-expressing A431 cell line and administered CATIPs intratumorally into this tumor. Because Nb_EGFR_ in CATIPs recognizes human, but not murine, EGFR, we simultaneously engrafted a second human EGFR-positive tumor, MDA-MB-468, at a separate site to model EGFR-expressing normal tissue. This dual-tumor design enabled us to assess whether molecules escaping from the treated tumor could induce OTOT toxicity in the distant, non-treated tumor, as well as systemic immune activation and exacerbation of xenogeneic graft-versus-host disease (xGVHD).^44^

**Figure 4.**
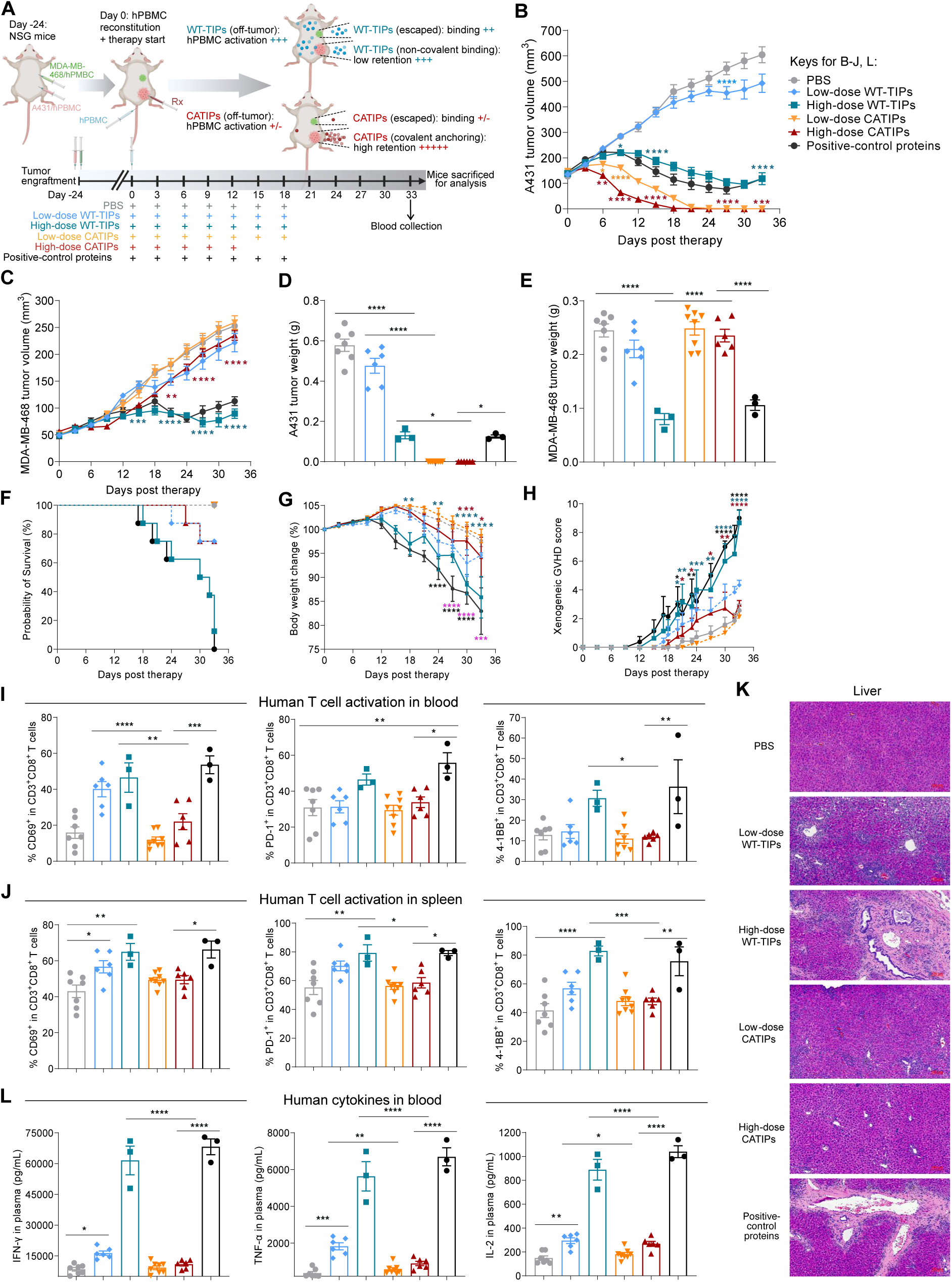
CATIP induced complete tumor clearance with limited toxicity in human PBMC-reconstituted NSG mice. **(A)** Experimental design for evaluating antitumor efficacy and OTOT toxicity after intratumoral administration of CATIPs or WT-TIPs in hPBMC-humanized NSG mice. **(B-C)** Growth curves of treated A431 tumors (**B**) and distant untreated MDA-MB-468 tumors (**C**). Symbols (see keys in panel B) are consistent across panels B–J and L. Data are mean ± SEM. n = 7 (PBS) and n = 8 (protein-treated groups) at study initiation. \**P* < 0.05, \*\**P* < 0.01, \*\*\**P* < 0.001, \*\*\*\**P* < 0.0001; asterisks colored as CATIPs denote comparisons with WT-TIPs at matched conditions, and asterisks colored as WT-TIPs denote comparisons with PBS (two-way ANOVA with Tukey’s multiple-comparisons test). **(D-E)** Endpoint weights of dissected A431 (**D**) or MDA-MB-468 (**E**) tumors. Data are mean ± SEM. n = 7 (PBS), 8 (low-dose CATIPs), 6 (low-dose WT-TIPs or high-dose CATIPs), or 3 (high-dose CATIPs or positive-control proteins). \**P* < 0.05, \*\*\*\**P* < 0.0001; one-way ANOVA with Tukey’s multiple-comparisons test. **(F)** Kaplan–Meier survival curves of mice following treatment. **(G)** Relative body-weight change during treatment. \**P* < 0.05, \*\**P* < 0.01, *** *P* < 0.001, \*\*\*\**P* < 0.0001; asterisks colored as CATIPs denote comparisons with WT-TIPs at matched conditions, asterisks colored as WT-TIPs or positive control denote comparisons with PBS, and pink asterisks denote comparisons between positive control proteins and high-dose CATIPs (two-way ANOVA with Tukey’s multiple-comparisons test). **(H)** xGVHD scores of mice following treatment. \**P* < 0.05, \*\**P* < 0.01, \*\*\**P* < 0.001, \*\*\*\**P* < 0.0001; asterisks colored as CATIPs denote comparisons with WT-TIPs at matched conditions, and asterisks colored as WT-TIPs or positive control denote comparisons with PBS (two-way ANOVA with Tukey’s multiple-comparisons test). **(I,J)** Activation of human T cells in blood (**I**) and spleen (**J**) of mice at study endpoint. \**P* < 0.05, \*\**P* < 0.01, \*\*\**P* < 0.001, \*\*\*\**P* < 0.0001; one-way ANOVA with Tukey’s multiple-comparisons test. (K) H&E staining of liver tissues to evaluate treatment-associated toxicity. (L) Human cytokine concentrations in blood of mice at study endpoint. \**P* < 0.05, \*\**P* < 0.01, \*\*\**P* < 0.001, \*\*\*\**P* < 0.0001; one-way ANOVA with Tukey’s multiple-comparisons test.

On day –24, A431 and MDA-MB-468 cells were each mixed with their respective partially HLA-matched PBMCs to limit allogeneic cytotoxicity during initial tumor establishment,^45^ and the mixtures were implanted subcutaneously at adjacent sites on the right flank, generating two neighboring tumors (**Fig. 4A**). By day 0, A431 and MDA-MB-468 tumors had reached approximately 150 mm³ and 50 mm³, respectively. hPBMCs partially HLA-matched to MDA-MB-468 cells were then adoptively transferred by tail-vein injection. On the same day, low or high doses of CATIPs or WT-TIPs were administered intratumorally into A431 tumors every 3 days. As a positive control, proteins comprising Nb_EGFR_-ScFv_UCHT1_, CD80, and Neo2/15, at molar amounts matched to the high-dose TIP combination, were delivered in the same manner. ScFv_UCHT1_, derived from the UCHT1 clone, is a well-established CD3 agonist.^46^

As expected, A431 tumors in PBS-treated mice grew rapidly (**Fig. 4B**). Low-dose WT-TIPs moderately delayed tumor growth after seven doses, whereas high-dose WT-TIPs and the positive-control proteins significantly suppressed A431 tumor growth after three doses and induced a sustained reduction in tumor volume. However, neither regimen eradicated A431 tumors, even after seven doses. By contrast, CATIPs at both dose levels showed markedly greater activity, inducing complete regression of A431 tumors after seven doses at the low dose and five doses at the high dose. Cleared tumors remained undetectable until the end of the study.

We next examined the non-treated MDA-MB-468 tumors as a surrogate for OTOT toxicity (**Fig. 4C**). High-dose WT-TIPs and the positive-control proteins significantly inhibited MDA-MB-468 tumor growth after four doses, and tumor volumes remained below 100 mm³ throughout the experiment. Low-dose WT-TIPs caused only transient growth arrest, followed by renewed tumor expansion. High-dose CATIPs produced an initial reduction in MDA-MB-468 tumor growth in two mice after three doses, but this effect was not significant, and growth subsequently resumed at a rate comparable to that in the PBS and low-dose CATIP groups. Notably, low-dose CATIPs had no measurable effect on MDA-MB-468 tumor growth, which remained comparable to that in the PBS-treated controls. Thus, unlike WT-TIPs, CATIPs retained potent activity against the injected A431 tumors while largely sparing the distant EGFR-positive MDA-MB-468 tumors.

Endpoint tumor weights were consistent with the longitudinal volume measurements. CATIPs at both dose levels completely eliminated A431 tumors, whereas WT-TIPs and the positive-control proteins reduced, but did not eliminate, tumor burden relative to PBS (**Fig. 4D**). In contrast, high-dose WT-TIPs and the positive-control proteins markedly reduced MDA-MB-468 tumor weight, whereas CATIPs had little effect relative to PBS (**Fig. 4E**).

The survival data further supported an improved safety profile for CATIPs (**Fig. 4F**). High-dose WT-TIPs and the positive-control proteins caused early mortality, with the first deaths occurring on days 18 and 17, respectively, and final mortality rates of 87.5% and 100%. In the high-dose CATIP and low-dose WT-TIP groups, the first deaths occurred later, on days 27 and 24, respectively, and mortality remained limited to 25% in each group. Notably, all mice in the PBS and low-dose CATIP groups survived to study endpoint.

Body weight and clinical xGVHD scores showed the same pattern. Mice in the PBS and low-dose CATIP groups lost weight only at late stages of the experiment (**Fig. 4G**), whereas mice treated with high-dose WT-TIPs or positive-control proteins exhibited earlier and more rapid weight loss, exceeding 15% by endpoint. Weight loss in the high-dose CATIP and low-dose WT-TIP groups emerged only during the middle to late phase of treatment. Clinical scoring likewise showed that high-dose WT-TIPs and the positive-control proteins markedly accelerated xGVHD progression relative to PBS and low-dose CATIPs (**Fig. 4H**), whereas high-dose CATIPs and low-dose WT-TIPs induced only mild xGVHD. Spleen atrophy at endpoint further reflected xGVHD severity in mice that succumbed earlier to treatment-associated toxicity (**Fig. S4A, S4B**).

To investigate the basis of this toxicity, we analyzed systemic T cell activation. In peripheral blood, both CD3⁺CD8⁺ and CD3⁺CD8⁻ T cells from mice treated with high-dose WT-TIPs or positive-control proteins increased expression of the activation markers CD69, PD-1, and 4-1BB (**Fig. 4I** **and S4C**). By contrast, T cells from CATIP-treated mice remained similar to those in PBS-treated controls. The same pattern was observed in splenic CD3⁺CD8⁺ and CD3⁺CD8⁻ T cells (**Fig. 4J** **and S4D**). Moreover, mice that died early after treatment with high-dose WT-TIPs or positive-control proteins showed markedly increased T cell activation at their terminal time points relative to mice receiving CATIPs or PBS, consistent with fatal xGVHD (**Fig. S5A-D**).

Consistent with these findings, high-dose WT-TIPs or positive-control proteins induced severe liver injury, a hallmark of hPBMC-driven xGVHD in this model. H&E staining revealed prominent lymphocyte-predominant portal inflammatory infiltrates and bridging fibrosis in groups treated with high-dose WT-TIPs or positive-control protein groups, whereas CATIP-treated mice showed histology comparable to PBS controls (**Fig. 4K**). WT-TIPs and positive-control proteins also increased circulating human cytokines, including IFN-γ, TNF-α, IL-2, IL-6, IL-10 and GM-CSF (**Fig. 4L** **and S6A**).^41–43^ By contrast, cytokine concentrations remained low in CATIP-treated mice and were similar to those in PBS controls. Mice that died early likewise showed markedly elevated cytokine levels at their terminal time points compared with CATIP-treated or PBS-treated mice at study endpoint, further supporting severe systemic immune activation as the cause of death (**Fig. S6B**).

These data show that CATIP combination therapy outperformed WT-TIPs and the matched positive-control proteins, completely eradicating the treated A431 tumors while sparing distant, untreated EGFR-positive MDA-MB-468 tumors and limiting systemic activation of adoptively transferred human PBMCs. CATIPs therefore combined superior local antitumor efficacy with reduced OTOT activity and xGVHD-associated toxicity, supporting their potential as a safer strategy for covalently anchored cancer immunotherapy.

### CATIPs induce antigen-specific abscopal effects and durable antitumor immune memory

Irreversible anchoring of CATIPs to tumor cells enables sustained, spatially confined delivery of immunostimulatory signals at the tumor site. We therefore asked whether intratumoral CATIP administration could remodel the local tumor microenvironment and elicit an abscopal response (**Fig. 5A**), whereby T cells primed in the treated tumor suppress distant, untreated tumors in immunocompetent mice.

**Figure 5.**
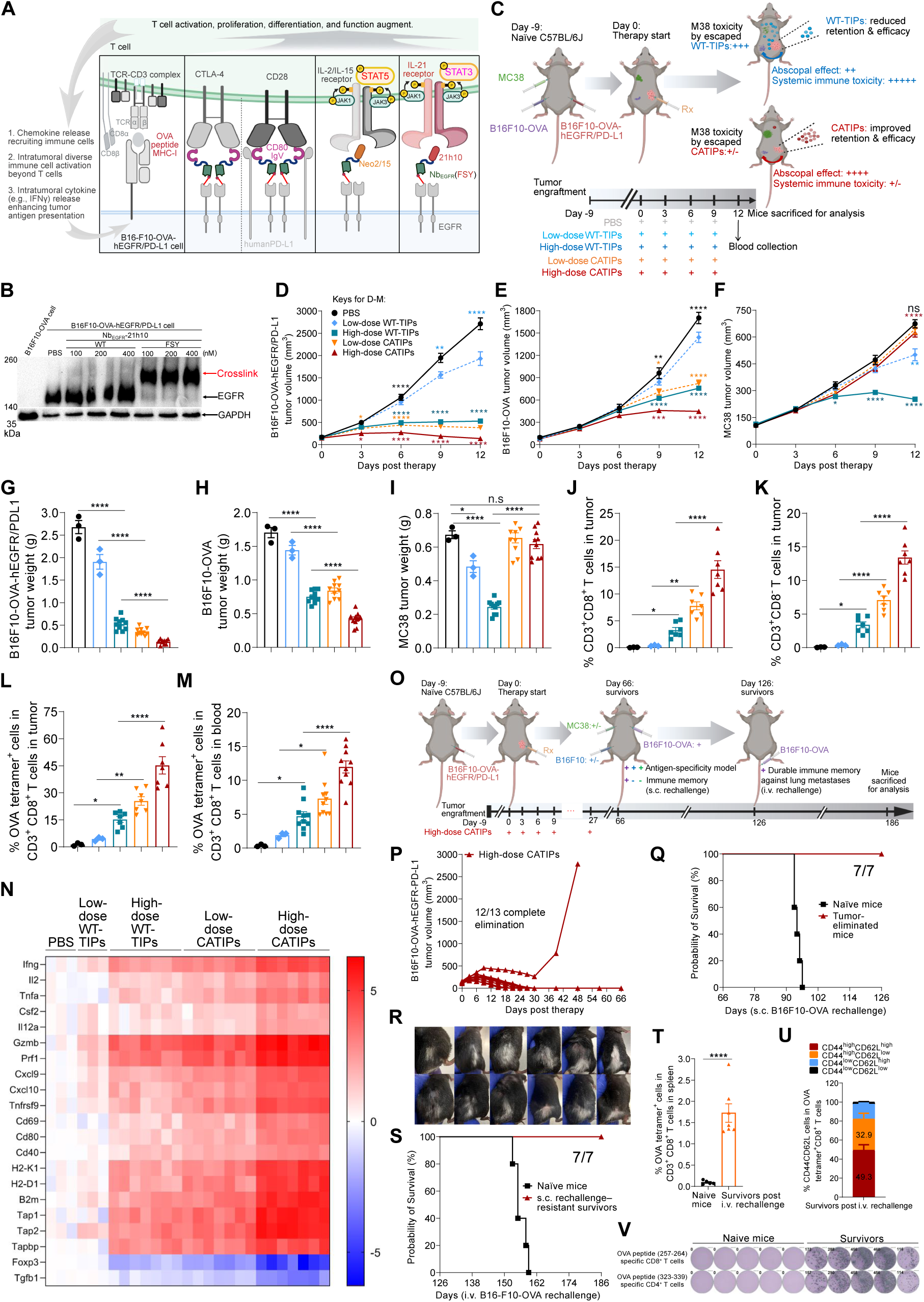
CATIPs induce local tumor regression, antigen-specific abscopal effects, and durable tumor-specific protective immunity in immunocompetent mice. (A) Schematic of CATIPs design for use in immunocompetent mice. Nb_EGFR_(FSY)–CD80, Nb_EGFR_(FSY)–Neo2/15, and Nb_EGFR_(FSY)–21h10 were used. (B) Nb_EGFR_(FSY)–21h10 covalently crosslinked human EGFR on the surface of B16F10-OVA-hEGFR/PD-L1 cells. (C) Experimental design for evaluating local antitumor efficacy, abscopal effects against distant untreated tumors, and specificity relative to antigen-unrelated tumors. **(D-F)** Growth curves of treated B16F10-OVA-hEGFR/PD-L1 tumors (**D**), distant untreated B16F10-OVA tumors (**E**), and antigen-unrelated MC38 tumors (**F**). Data are mean ± SEM. n = 10 (PBS) or 13 (TIPs-treated groups) at study initiation. n.s. *P* > 0.05; \**P* < 0.05, \*\**P* < 0.01, \*\*\**P* < 0.001, \*\*\*\**P* < 0.0001; asterisks colored as CATIPs denote comparisons with WT-TIPs at matched conditions, asterisks colored as WT-TIPs denote comparisons with PBS, and black asterisks denote comparisons between low-dose CATIPs and PBS (two-way ANOVA with Tukey’s multiple-comparisons test). **(G-I)** Endpoint weights of treated B16F10-OVA-hEGFR/PD-L1 tumors (**G**), distant untreated B16F10-OVA tumors (**H**), and antigen-unrelated MC38 tumors (**I**). Data are mean ± SEM. n = 3 (PBS or low-dose WT-TIPs) or 10 (high-dose WT-TIPs or CATIP-treated groups) at study endpoint. n.s. *P* > 0.05, \**P* < 0.05, \*\*\*\**P* < 0.0001; one-way ANOVA with Tukey’s multiple-comparisons test. **(J,K)** Frequencies of intratumoral CD3^+^ CD8^+^ T cells (**J**) and CD3^+^ CD8^-^ T cells (**K**). \**P* < 0.05, \*\**P* < 0.01, \*\*\*\**P* < 0.0001; one-way ANOVA with Tukey’s multiple-comparisons test. **(L,M)** Frequencies of OVA tetramer^+^ cells among CD3^+^ CD8^+^ T cells in tumors (**L**) and blood (**M**). \**P* < 0.05, \*\**P* < 0.01, \*\*\*\**P* < 0.0001; one-way ANOVA with Tukey’s multiple-comparisons test. **(N)** Heatmap of immune-related gene expression in bulk tum or RNA measured by qPCR. **(O)** Experimental scheme for evaluation of tumor eradication and subsequent immune memory after rechallenge. **(P)** Individual growth curves of high-dose CATIP-treated B16F10-OVA-hEGFR/PD-L1 tumors. **(Q)** Kaplan–Meier survival after subcutaneous rechallenge with B16F10-OVA cells to assess immune memory. n = 5 (naïve mice) and n = 7 (tumor-eliminated mice). **(R)** Representative images showing vitiligo-like depigmentation and local hair whitening at day 96. **(S)** Kaplan–Meier survival after intravenous rechallenge with B16F10-OVA cells to assess protection against lung metastasis. n = 5 (naïve mice) and n = 7 (survivors resistant to prior subcutaneous rechallenge). **(T)** Frequency of OVA tetramer^+^ cells among splenic CD3^+^ CD8^+^ T cells at day 186. Data are mean ± SEM. n = 5 (naïve mice) and n = 7 (survivors after i.v. rechallenge). \*\*\*\**P* < 0.0001; unpaired *t*-test. **(U)** Frequencies of CD44^high^CD62L^high^ central memory and CD44^high^CD62L^low^ effector memory subsets among splenic OVA tetramer^+^ CD8^+^ T cells at day 186. **(V)** Representative IFN-γ ELISpot images and quantification of splenocytes following recall stimulation with OVA peptides. Numbers denote spot-forming units per well, representing individual IFN-γ-secreting cells.

To test this in an immunocompetent syngeneic setting, the immunomodulatory payloads needed to engage murine immune receptors. We used B16F10-OVA, a B16F10-derived murine melanoma line stably expressing chicken ovalbumin (OVA), as a model antigen. To generate a primary tumor suitable for CATIP anchoring, we established B16F10-OVA-hEGFR/PD-L1 cells stably expressing human EGFR for CATIP attachment and human PD-L1 to support activity of the CD80 IgV variant (**Fig. S7A-D**), which recognize human but not mouse PD-L1 and activates mouse CD28 in a human PD-L1 dependent manner while simultaneously blocking mouse CTLA-4. Neo2/15 engages both human and mouse receptors.^39^ Nb_EGFR_(FSY)-CD80 and Nb_EGFR_(FSY)-Neo2/15 also efficiently crosslinked B16F10-OVA-hEGFR/PD-L1 cells (**Fig. S7E-F**).

Because the human CD3ε agonistic nanobody Nb_CD3_ is inactive on murine CD3ε, we replaced the CD3 module with IL-21 (**Fig. 5A**), a potent immunostimulatory cytokine that enhances CD8 T cell and NK cell function and promotes memory formation, while reducing regulatory T cell activity, but is associated with substantial systemic toxicity.^47,48^ These properties made IL-21 well suited for spatially restricted, covalently anchored delivery. We accordingly generated Nb_EGFR_(FSY)-21h10, a CATIP carrying the potent IL-21 mimic 21h10,^48^ and confirmed efficient covalent crosslinking to EGFR ECD in vitro (**Fig. S7G**) and to native EGFR on B16F10-OVA-hEGFR/PD-L1 cells (**Fig. 5B**) and MDA-MB-486 cells (**Fig. S7H**). 21h10 activates STAT3 signaling whereas Neo2/15 activates STAT5 signaling, potentially enabling complementary immunostimulatory effects.^49^

We next assessed the activity of these CATIPs on murine immune cells by co-culturing naïve C57BL/6 splenocytes with B16F10-OVA-hEGFR/PD-L1 cells in the presence of CATIPs. Under non-wash conditions, CATIP combinations enhanced splenocyte–mediated tumor cell killing relative to the corresponding WT-TIP combinations (**Fig. S7I**). After PBS washing, WT-TIP activity was largely lost, whereas CATIPs retained robust cytotoxicity against B16F10-OVA-hEGFR/PD-L1 cells (**Fig. S7J**).

We then tested whether intratumoral administration of these CATIPs could suppress or regress the treated tumors in situ and induce an abscopal effect against distant untreated tumors while sparing antigen-unrelated tumors. Immunocompetent C57BL/6 mice were inoculated subcutaneously with B16F10-OVA–hEGFR/PD-L1 cells as the primary tumor, B16F10-OVA cells as a distant untreated antigen-matched tumor, and MC38 cells as a distant untreated antigen-unrelated tumor (**Fig. 5C**).^50^ When primary B16F10-OVA–hEGFR/PD-L1 tumors reached 150–200 mm³, WT-TIPs or CATIPs were administered intratumorally every 3 days at low or high dose.

Primary B16F10-OVA–hEGFR/PD-L1 tumors in PBS-treated mice grew rapidly (**Fig. 5D and Fig. S8A**). Three low-dose WT-TIP treatments slowed tumor growth, whereas two doses of high-dose WT-TIPs or low-dose CATIPs significantly inhibited tumor growth relative to PBS. High-dose CATIPs were markedly more active than high-dose WT-TIPs, with a clear effect after the first dose. After two high-dose CATIP treatments, tumor growth was completely arrested, and after four doses tumors showed modest regression toward their pretreatment size. By day 12, low-dose CATIPs had also induced tumor regression, whereas high-dose WT-TIPs produced growth inhibition without significant regression. Thus, CATIPs showed greater local antitumor efficacy than WT-TIPs at matched doses.

We next assessed the abscopal response in distant untreated B16F10-OVA tumors. High-dose CATIPs significantly slowed tumor growth after three doses and completely arrested growth after four doses (**Fig. 5E and Fig. S8B**). By contrast, three doses of high-dose WT-TIPs or low-dose CATIPs only delayed tumor growth relative to PBS and did not fully suppress tumor expansion after four doses. Thus, high-dose CATIPs elicited a stronger abscopal effect against distant antigen-matched tumors than WT-TIPs, despite the ability of WT-TIPs to diffuse away from the injected tumor.

The response of the antigen-unrelated MC38 tumors was distinct. High-dose WT-TIPs significantly suppressed MC38 tumor growth after three doses and induced regression after four doses (**Fig. 5F and Fig. S8C**). In contrast, CATIPs at either high or low dose did not significantly affect MC38 growth. These findings indicate that CATIPs induced an antigen-restricted abscopal effect against B16F10-OVA tumors without affecting antigen-unrelated MC38 tumors, despite the MC38 model being immunologically “hot”.^38,48^ By contrast, WT-TIPs appeared to disseminate systemically and activate broader peripheral immunity, thereby suppressing MC38 growth independently of EGFR binding, as MC38 cells do not express the anchoring target.

Endpoint tumor weights were consistent with the longitudinal growth data. CATIPs at both dose levels significantly reduced the weight of primary B16F10-OVA–hEGFR/PD-L1 tumors (**Fig. 5G**). Although high-dose WT-TIPs also reduced tumor weight relative to PBS, the mean tumor mass remained approximately 4-fold higher than in the high-dose CATIP group. Likewise, mean tumor mass in the low-dose WT-TIP group was approximately 5.4-fold higher than in the low-dose CATIP group. CATIPs therefore reduced local tumor burden substantially more effectively than WT-TIPs.

CATIPs also significantly reduced the weight of distant untreated B16F10-OVA tumors at both doses (**Fig. 5H**). High-dose WT-TIPs also reduced tumor weight relative to PBS, but the mean tumor mass remained approximately 1.8-fold higher than in the high-dose CATIP group. The corresponding difference at low dose was approximately 1.7-fold. In contrast, high-dose WT-TIPs reduced MC38 tumor weight to approximately 39.5% of that observed in high-dose CATIP–treated mice (**Fig. 5I**), further underscoring the distinct off-tumor profiles of WT-TIPs and CATIPs.

To investigate the basis of these differences, we analyzed tumor-infiltrating lymphocytes. High-dose CATIPs significantly increased the frequency of intratumoral CD3⁺CD8⁺ T cells (**Fig. 5J**) and CD3⁺CD8⁻ T cells (**Fig. 5K**) relative to all other groups, reaching approximately 4.5-fold and 4-fold higher levels, respectively, than in the high-dose WT-TIP group. High-dose CATIPs also markedly expanded OVA-specific CD3⁺CD8⁺ T cells within tumors, to approximately 3-fold the levels observed with high-dose WT-TIPs (**Fig. 5L and S8D**), consistent with previous reports that OVA-specific T cells can be induced and expanded in wild-type C57BL/6 mice.^51^ Even low-dose CATIPs significantly increased intratumoral OVA-specific CD3⁺CD8⁺ T cells, to approximately 5.9-fold the level induced by low-dose WT-TIPs (**Fig. 5L**). Similarly, high-dose and low-dose CATIPs increased OVA-specific CD3⁺CD8⁺ T cells in peripheral blood by approximately 2.5-fold and 3.8-fold, respectively, relative to WT-TIPs (**Fig. 5M and S8E**). Expansion of circulating OVA-specific CD3⁺CD8⁺ T cells probably contributed to the observed abscopal control of distant B16F10-OVA tumors.

qPCR analysis of bulk tumor RNA showed that CATIP treatment induced broad immune activation within the tumor bed, increasing expression of cytokines (IFNγ, TNFα, IL-2, IL-12a, and GM-CSF), effector molecules (GZMB and PRF1), chemokines (CXCL9 and CXCL10), T-cell activation markers (4-1BB and CD69), antigen-presenting cell activation markers (CD80 and CD40), and antigen-presentation machinery components (H2K1, H2D1, B2M, TAP1, TAP2, and Tapbp)^3^ relative to other treatments. At the same time, CATIP treatment reduced expression of the immunosuppressive factor TGF-β and the Treg-transcription factor Foxp3 (**Fig. 5N**). These data indicate that intratumoral CATIP administration induced a strong inflammatory program associated with enhanced antigen presentation, increased effector differentiation, and reduced local immune suppression. This transcriptional remodeling is consistent with conversion of the treated tumor into a highly immunostimulatory niche that supports priming and expansion of tumor-reactive T cells.^52,53^

We next asked whether repeated high-dose CATIP treatment could eradicate established tumors, generate durable protective immunity, and do so without overt systemic toxicity. In a parallel experiment, C57BL/6 mice bearing only B16F10-OVA–hEGFR/PD-L1 tumors received high-dose CATIPs intratumorally every 3 days (**Fig. 5O**). High-dose CATIP treatment completely eradicated established tumors in 12 of 13 mice (92.3%) after 6 to 10 doses (**Fig. 5P**). Repeated CATIP administration did not decrease body weight, suggesting a lack of overt systemic toxicity (**Fig. S9A**). No tumor recurrence was observed during follow-up in mice that achieved complete regression.

To assess the specificity of the resulting immune protection, tumor-free survivors were rechallenged subcutaneously on day 66 with B16F10-OVA, B16F10 or MC38 cells (**Fig. 5O**). B16F10-OVA and B16F10 were used to test protection against melanoma, whereas MC38 served as an antigen-unrelated control. Mice that had previously cleared tumors rejected both B16F10-OVA and B16F10 rechallenge, with no detectable tumor growth through day 96, whereas naïve mice showed rapid tumor progression **(Fig. S9B-C)**. By contrast, MC38 tumors grew similarly in tumor-cleared survivors and naïve controls **(Fig. S9D)**. These data indicate that CATIP treatment induced durable, antigen-specific protective immunity against B16F10 melanoma, including OVA-negative B16F10 cells, without conferring nonspecific resistance to unrelated MC38 tumors.

In a parallel cohort, tumor-free survivors were rechallenged with B16F10-OVA cells alone and monitored for long-term survival (**Fig. 5O**). All rechallenged survivors remained tumor-free and survived throughout the study, whereas naïve mice developed rapidly progressing tumors and reached 100% mortality by day 96 (**Fig. 5Q**).

Interestingly, mice that achieved complete tumor regression developed vitiligo-like depigmentation with local hair whitening confined to the treated area (**Fig. 5R**), a phenotype consistent with immune responses against shared melanocytic antigens, as reported in preclinical melanoma models and in patients responding to immunotherapy.^31,54,55^

To further test whether systemic protection extended to metastatic disease, we intravenously injected B16F10-OVA cells on day 126, exploiting the lung-colonizing capacity of these cells, into survivors that had previously undergone subcutaneous rechallenge with B16F10-OVA cells alone (**Fig. 5O**). Strikingly, these mice remained free of metastatic lung foci, whereas naïve mice developed extensive pulmonary metastases (**Fig. S9E**). Accordingly, rechallenge-resistant survivors showed durable tumor-free survival to the end of study (day 186), while naïve control mice all developed lung metastases and died by days 153-159 (**Fig. 5S**). Flow cytometry analysis of splenocytes showed that OVA-specific CD3⁺CD8⁺ T cells were still detectable in lung metastasis–resistant survivor mice on day 186 (**Fig. 5T** and **Fig. S9F**). Among these OVA-specific CD3⁺CD8⁺ T cells, 49.3% were CD44^high^CD62L^high^ central memory T cells and 32.9% were CD44^high^CD62L^low^ effector memory T cells (**Fig. 5U** and **Fig. S9G**). Upon OVA peptide stimulation, splenocytes secreted IFN-γ, as detected by ELISpot (**Fig. 5V**), indicating that both OVA-specific CD8⁺ and CD4^+^ T cells retained antigen-specific recall responses upon re-exposure. Three of seven survivors developed vitiligo-like depigmentation with localized hair whitening at the subcutaneous B16F10-OVA rechallenge site (**Fig. S9H**). In addition, all survivors (7/7) maintained stable vitiligo-like depigmentation with localized hair whitening at the initial treatment site throughout the treatment period, without further expansion of the depigmented area. No detectable off-tumor toxicity was observed in tissues from CATIP-treated survivors relative to naïve controls, and no histological evidence of pancreas injury—a toxicity previously reported for 21h10^48^—was identified (**Fig. S9I**).

Intratumorally administered CATIPs therefore outperformed WT-TIPs by producing stronger local tumor control, eliciting a robust antigen-specific abscopal effect, and establishing durable protective immune memory. Repeated CATIP treatment eradicated established primary tumors, protected against both subcutaneous and intravenous rechallenge, and did so without detectable systemic toxicity.

### CATIPs–engineered whole-tumor-cell vaccines induce durable tumor-specific protective immunity

The irreversible anchoring of CATIPs to tumor-associated antigens suggested a distinct vaccine strategy. We reasoned that pre-treating B16F10-OVA–hEGFR/PD-L1 cells with CATIPs before implantation would generate engineered whole-tumor-cell vaccines that retain covalently bound immunostimulatory proteins on the cell surface, thereby converting an otherwise poorly immunogenic tumor cell into an antigen-presenting-cell-like vaccine that presents co-stimulatory and cytokine cues at the tumor cell–T cell interface (**Fig. 6A**).^56^ By contrast, analogous WT-TIPs were expected to dissociate and therefore provide insufficient and unstable stimulation in vivo.

**Figure 6.**
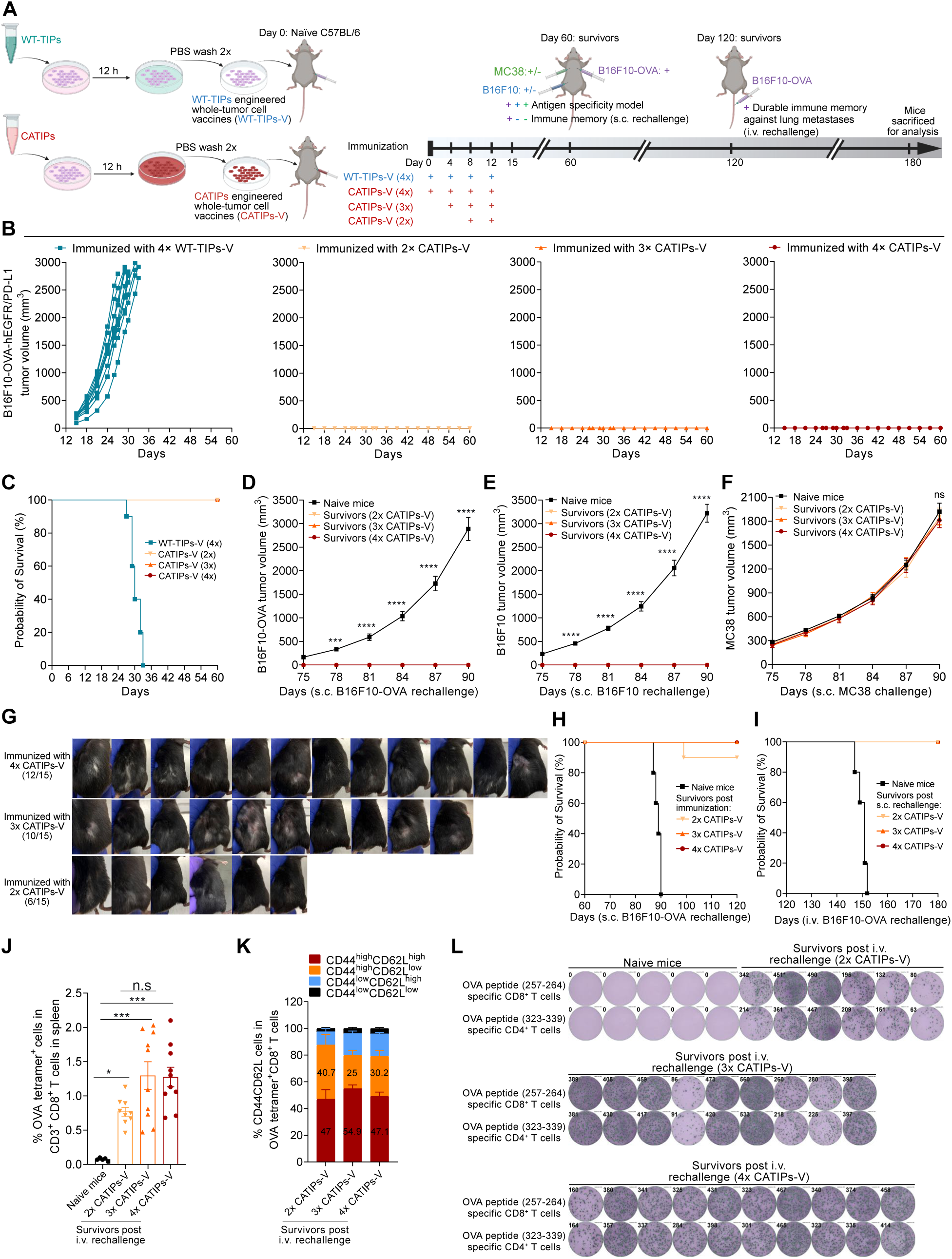
CATIP–engineered whole-tumor-cell vaccines induce durable tumor-specific protective immunity. **(A)** Schematic of the generation and vaccination strategy for CATIP-engineered whole-tumor-cell vaccines. **(B)** Individual growth curves of implanted TIP-engineered vaccine cells. n = 10 (WT-TIPs-V) and n = 15 (CATIPs-V). **(C)** Kaplan–Meier survival of vaccinated mice. **(D-F)** Growth curves after subcutaneous rechallenge with B16F10-OVA (**D**), B16F10 (**E**), or MC38 (**F**). Data are mean ± SEM. n = 5 mice. n.s. *P* > 0.05; \**P* < 0.05, \*\**P* < 0.01, \*\*\*\**P* < 0.0001; two-way ANOVA with Tukey’s multiple-comparisons test. **(G)** Images of mice showing vitiligo-like depigmentation and local hair whitening after vaccination at day 90. **(H)** Kaplan–Meier survival following subcutaneous rechallenge with B16-F10-OVA cells in mice previously vaccinated with CATIPs-V. n = 5 (naïve mice) and n = 10 (CATIPs-V vaccinated mice). **(I)** Kaplan–Meier survival following intravenous rechallenge with B16-F10-OVA cells in mice previously vaccinated with CATIPs-V and resistant to prior subcutaneous challenge. n =5 (naïve mice), n = 9 (2x CATIPs-V), and n = 10 (3x or 4x CATIPs-V). **(J)** Frequency of OVA tetramer^+^ cells among splenic CD3^+^ CD8^+^ T cells at day 180. Mice were vaccinated with CATIPs-V, followed by subcutaneous and intravenous rechallenge. n = 5 (naïve mice), n = 9 (2× CATIPs-V), and n = 10 (3× or 4× CATIPs-V). n.s., *P* > 0.05; \**P* < 0.05; \*\*\**P* < 0.001 (one-way ANOVA with Tukey’s multiple-comparisons test). **(K)** Frequencies of CD44^high^CD62L^high^ central memory and CD44^high^CD62L^low^ effector memory subsets among splenic OVA tetramer^+^ CD8^+^ T cells at day 180. **(L)** Representative IFN-γ ELISpot images and quantification of splenocytes following recall stimulation with OVA peptides. Numbers denote spot-forming units per well, representing individual IFN-γ-secreting cells.

To test this concept, we immunized C57BL/6 mice with CATIP–engineered whole-tumor-cell vaccines administered for two, three, or four rounds; mice receiving four rounds of WT-TIP–engineered tumor cells served as controls (**Fig. 6A**). WT-TIP–engineered cells formed rapidly growing tumors, whereas mice receiving two, three, or four rounds of CATIP–engineered vaccines showed no detectable tumor growth (**Fig. 6B**). Accordingly, all mice in the WT-TIP group died by day 33, whereas mice immunized with CATIP–engineered vaccines remained tumor-free and survived through day 60 (**Fig. 6C)**. Thus, covalent surface engineering of CATIPs rendered these tumor cells sufficiently immunogenic to prevent progressive outgrowth after implantation.

We next asked whether vaccination generated protective immunity. On day 60, surviving mice were divided into two cohorts and rechallenged subcutaneously (**Fig. 6A**). Consistent with the results in **Fig. S9B-D**, vaccinated survivors rejected both B16F10-OVA and B16F10 rechallenge, with no detectable tumor growth during follow-up, whereas naïve mice showed rapid tumor progression (**Fig. 6D-E**). By contrast, MC38 tumors grew similarly in vaccinated and naïve mice (**Fig. 6F**). These data indicate that CATIP-engineered tumor-cell vaccination induced tumor-specific protective immunity against both OVA-positive and OVA-negative B16F10 melanoma, without affecting the growth of an antigen-unrelated tumor.

In a parallel cohort, vaccinated survivors were rechallenged subcutaneously with B16F10-OVA alone and monitored for long-term survival. Mice previously immunized with CATIP-engineered vaccines remained tumor-free, with 100% survival in the three-dose and four-dose groups and 90% survival in the two-dose group through day 120, whereas naïve mice developed rapidly progressive tumors and reached 100% mortality by day 90 (**Fig. 6H**).

As in mice that cleared established tumors after intratumoral CATIP therapy (**Fig. 5R**), mice immunized with CATIP–engineered vaccines developed vitiligo-like depigmentation with local hair whitening at the vaccination site (**Fig. 6G**). The frequency of this phenotype increased with the rounds of immunizations, consistent with progressively stronger immune responses against shared melanocytic antigens.

To determine whether this protection extended to metastatic challenge, mice that had rejected subcutaneous rechallenge were injected intravenously with B16F10-OVA cells on day 120 (**Fig. 6A**). Survivors resistant to subcutaneous B16F10-OVA rechallenge remained free of metastatic lung foci in all mice vaccinated with three or four rounds of CATIP-engineered vaccines (10/10), and in 7 of 9 mice vaccinated with two rounds of CATIP-engineered vaccines, whereas naïve mice developed extensive pulmonary metastases (**Fig. S10A**). Correspondingly, rechallenge-resistant survivors showed durable tumor-free survival to the end of study (day 180), while naïve controls all died by days 147-152 (**Fig. 6I**).

Flow cytometric analysis of splenocytes from survivors showed presence of OVA-specific CD8⁺ T cells on day 180 (**Fig. 6J and Fig. S10B**). Among these cells, CD44^high^CD62L^high^ central memory and CD44^high^CD62L^low^ effector memory subsets were identified, with both subsets present at substantial proportions (**Fig. 6K and Fig. S10C**). Consistent with **Fig. 5V**, collected splenocytes produced IFN-γ upon re-stimulation with OVA peptides, as measured by ELISpot (**Fig. 6L**), indicating that both OVA-specific CD8⁺ and CD4^+^ T cells retained functional recall responses upon antigen re-exposure. These findings indicate maintenance of B16F10-OVA–reactive effector and memory T cells after vaccination and rechallenge. In addition, all survivors maintained stable depigmentation confined to the initial vaccination site without further expansion, and a subset also developed vitiligo-like depigmentation with localized hair whitening at the subcutaneous B16F10-OVA rechallenge site (**Fig. S10D**).

These results show that covalent surface engineering with CATIPs converted poorly immunogenic tumor cells into effective whole-cell vaccines that did not establish tumors but instead elicited durable, tumor-specific immune protection. Vaccinated mice rejected both subcutaneous and intravenous tumor rechallenge, remained protected from lung metastasis, and generated functional tumor-reactive memory T cell responses.

### A CATIP library expands local immune-targeting strategies

The modularity of covalent anchoring prompted us to ask whether this platform could be extended beyond the CATIPs described above to generate a broader set of tumor-retained immunotherapies. To enable local combination therapy across multiple immune compartments, we developed a covalently anchored library designed to block suppressive pathways and engage T cells, NK cells, NKT cells, γδ T cells and myeloid cells in situ.

To localize checkpoint blockade, we constructed covalently anchored inhibitors targeting PD-1, PD-L1, CTLA-4, TIM-3, LAG-3 and TIGIT (**Fig. 7A**).^7^ Each fusion protein efficiently crosslinked EGFR on the surface of A431 cells (**Fig. 7B-G**), establishing a panel of tumor-retained checkpoint inhibitors that could, in principle, both block inhibitory signaling and engage checkpoint-expressing tumor-infiltrating immune cells.

**Figure 7.**
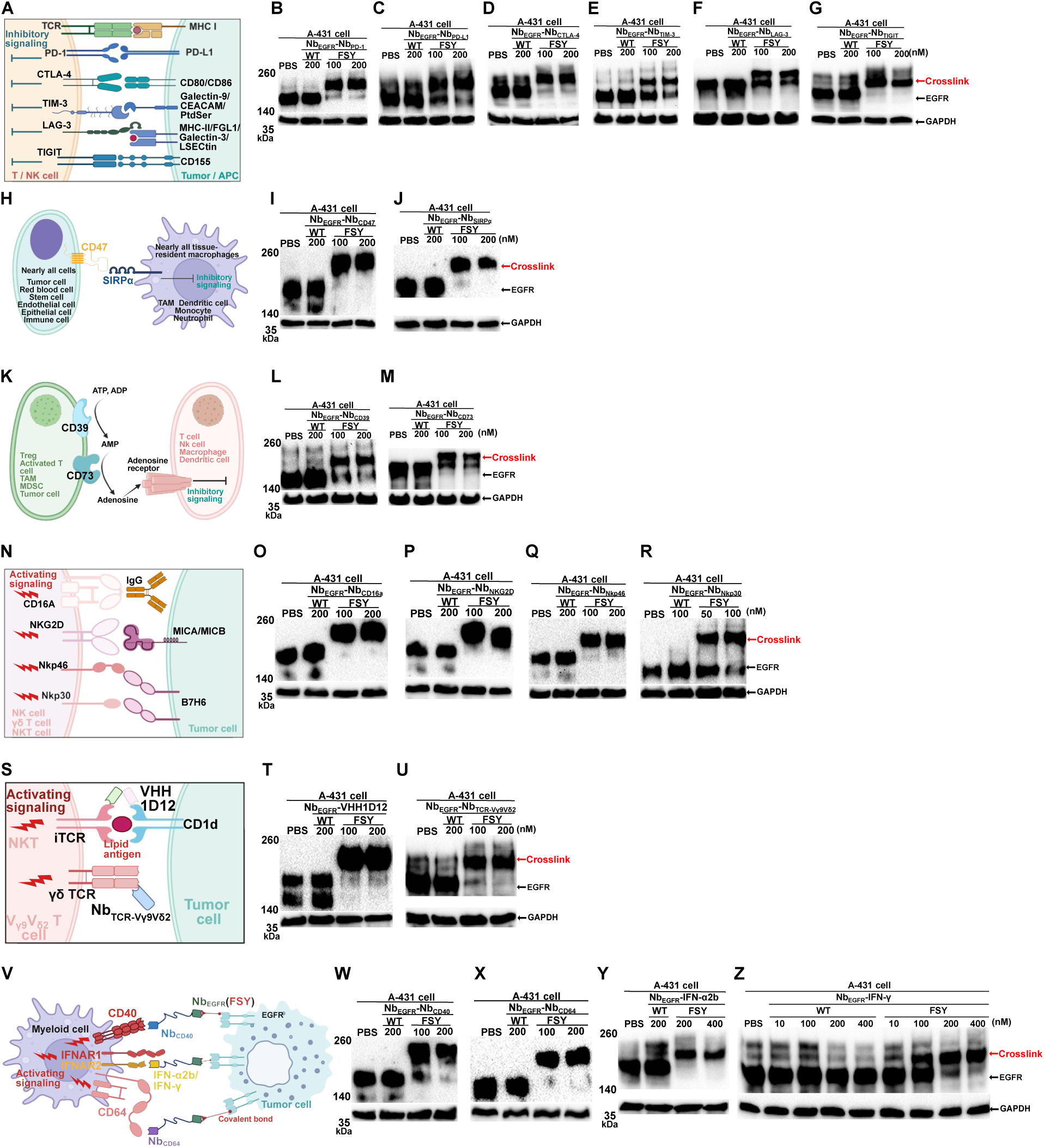
Covalently anchored protein modules expand the CATIP platform. **(A)** Schematic of classical immune checkpoints (ICs). PtdSer: Phosphatidylserine. **(B-G)** Nb_EGFR_(FSY)-Nb_IC_ constructs covalently crosslinked EGFR on the surface of A431 cells. **(H)** Schematic of classical phagocytosis checkpoints (PCs). **(I,J)** Nb_EGFR_(FSY)-Nb_PC_ constructs covalently crosslinked EGFR on the surface of A431 cells. **(K)** Schematic of classical immunometabolic checkpoints (IMCs). **(L,M)** Nb_EGFR_(FSY)-Nb_IMC_ constructs covalently crosslinked EGFR on the surface of A431 cells. **(N)** Schematic of classical NK cell-activating receptors (NKARs). **(O-R)** Nb_EGFR_(FSY)-Nb_NKAR_ constructs covalently crosslinked EGFR on the surface of A431 cells. **(S)** Schematic of classical NKT- and γδ T cell-activating receptors. **(T,U)** Nb_EGFR_(FSY)-VHH1D12 (**T**) or Nb_EGFR_(FSY)-Nb_TCR-Vγ9Vδ2_ (**U**) covalently crosslinked EGFR on the surface of A431 cells. **(V)**Schematic of classical myeloid cell-activating receptors (MCARs). **(X-Z)** Nb_EGFR_(FSY)-Nb_MCAR_ constructs covalently crosslinked EGFR on the surface of A431 cells.

We next extended this strategy to additional suppressive pathways in the tumor microenvironment. To localize blockade of the phagocytosis checkpoint (**Fig. 7H**),^57^ we generated Nb_EGFR_(FSY)–Nb_CD47_ and Nb_EGFR_(FSY)–Nb_SIRPα_, both of which efficiently crosslinked EGFR on A431 cells (**Fig.7I-J**). We similarly targeted the ATP–adenosine axis by generating Nb_EGFR_(FSY)–Nb_CD39_ and Nb_EGFR_(FSY)–Nb_CD73_ (**Fig. 7K**),^58^ and both constructs showed efficient covalent tethering on A431 cells (**Fig. 7L-M**).

To broaden immune cell-engagement beyond conventional T cells, we generated a panel of covalently anchored NK cell-stimulatory proteins, including Nb_EGFR_(FSY)–Nb_CD16_, Nb_EGFR_(FSY)–Nb_NKG2D_, Nb_EGFR_(FSY)–Nb_NKp46_, and Nb_EGFR_(FSY)–Nb_NKp30_ (**Fig. 7N**).^59^ All four constructs efficiently crosslinked EGFR on A431 cells (**Fig. 7O-R**). We also generated Nb_EGFR_(FSY)–VHH1D12 to engage type I NKT cells and Nb_EGFR_–Nb_TCR-Vγ9Vδ2_ to target Vγ9Vδ2 T cells (**Fig. 7S**),^60,61^ and both proteins showed efficient covalent tethering (**Fig. 7T-U**). These constructs expand the platform to innate-like lymphocyte populations that can mediate antitumor activity independently of classical peptide-MHC recognition, including Vγ9Vδ2 T cells, which are emerging as promising immunotherapy effectors.^62^

Finally, to target myeloid cells, we generated Nb_EGFR_(FSY)–Nb_CD40,_ Nb_EGFR_(FSY)–Nb_CD64_, Nb_EGFR_(FSY)–IFNα2b, and Nb_EGFR_(FSY)–IFNγ proteins (**Fig. 7V**).^63^ Each construct exhibited efficient covalent membrane tethering (**Fig. 7W-Z**). This establishes a versatile library of covalently anchored, tumor-retained immune modulators spanning checkpoint blockade, innate immune activation, and myeloid reprogramming, and provides a framework for rational local combination immunotherapy.

## DISCUSSION

This study establishes covalent tumor anchoring that spatially organizes immunotherapy at the tumor-cell surface, converting diffusible immunostimulatory proteins into durable, membrane-confined immune cues that uncouple local potency from systemic exposure. Across human PBMC-reconstituted NSG mice and immunocompetent syngeneic melanoma models, CATIPs outperformed matched non-covalent proteins, eradicating established treated tumors while limiting systemic T cell activation, cytokine release, on-target, off-tumor toxicity, and treatment-related morbidity. In immunocompetent settings, local CATIP therapy induced antigen-specific responses in distant tumors and established durable protective memory. Consistent with this, CATIP-engineered tumor cells functioned as whole-cell vaccines that protected against subsequent local and metastatic challenge, indicating that covalent surface engineering can convert poorly immunogenic tumor cells into immunostimulatory platforms for tumor-specific protection.

Mechanistically, covalent anchoring fundamentally reconfigures the spatiotemporal organization of immune signaling at tumor-cell surfaces. Soluble T cell engagers rely on reversible trimolecular interactions among the engager, the tumor antigen and the TCR-CD3 complex, processes that are intrinsically constrained by dissociation, diffusion and non-productive receptor occupancy.^64,65^ By contrast, CATIPs convert this transient interaction into a stable, membrane-confined signaling interface at the tumor-cell surface. Once anchored, activating ligands persist on the tumor surface, enabling sustained, synapse-like stimulation upon immune-cell contact. This framework helps explain the enhanced potency of CATIPs and their capacity to drive complete tumor eradication across experimental settings. It may also help address limitations of soluble T cell engagers, in which high exposure can promote non-productive receptor engagement.^18^

Covalent tumor anchoring functionally recapitulates key features of physiological T cell activation in a tumor-restricted format. The EGFR-targeted CATIP combination delivered CD3 engagement for signal 1, CD80-mediated CD28 co-stimulation for signal 2, and IL-2/IL-15 receptor agonism for signal 3. These inputs approximate the canonical signals delivered by antigen-presenting cells (APCs),^66^ yet are imposed directly on tumor cells without TCR reprogramming or genetic manipulation. Rather than converting tumor cells into professional APCs, CATIPs establish an APC-like interface on malignant cells, enabling localized immune activation while minimizing systemic engagement.

A key consequence of this design is spatial control. Previous approaches to localize cytokines or immune agonists have shown that spatial confinement can improve therapeutic index,^15,17^ but these interactions remain reversible. Covalent anchoring instead retains immunostimulatory proteins at the tumor cell surface, sustaining local signaling while limiting systemic redistribution. In vivo, non-covalent WT-TIPs redistributed into the circulation and suppressed distant EGFR-positive tumors, whereas CATIPs remained confined to the injected tumor and largely spared untreated EGFR-positive tumors. This difference translated into a substantially improved therapeutic window, enabling complete tumor eradication while limiting systemic T cell activation, inflammatory cytokine release and tissue toxicity.

CATIPs elicited antigen-specific systemic immunity, demonstrating that covalent tumor anchoring preserves the capacity of local therapy to generate systemic antitumor responses. In the melanoma model, CATIPs controlled distant antigen-matched tumors but not antigen-unrelated tumors, whereas WT-TIPs suppressed both, consistent with broader systemic distribution. These findings indicate that covalent anchoring maintains tumor-antigen specificity while permitting systemic immune propagation. Beyond antigen specificity, CATIPs induced coordinated remodeling of the tumor immune microenvironment. Intratumoral CATIP administration increased tumor infiltration by CD8⁺ and CD8⁻ T cells, expanded antigen-specific CD8⁺ T cells in tumors and blood, and induced intratumoral transcriptional programs associated with inflammation, T cell activation, cytotoxic effector function, antigen-presentation machinery and endogenous APC activation, while reducing immunosuppressive mediators such as TGF-β and Foxp3.^67^ These changes indicate that CATIPs shift the tumor microenvironment from an immunosuppressive state toward a T cell-inflamed, antigen-presenting niche capable of priming and expanding tumor-reactive T cells.

The durability of this response suggests that CATIP-mediated tumor destruction promotes antigen spreading and memory formation. Mice that cleared established tumors rejected subsequent tumor rechallenge, resisted metastatic challenge and retained functional central and effector memory CD8⁺ T cells. The vitiligo-like depigmentation observed in tumor-cleared mice is consistent with immune recognition of endogenous tumor-associated antigens beyond the engineered model antigen.^31,54,55,68^ Together, these findings indicate that localized tumor destruction can initiate systemic, antigen-diversified immunity with long-term protective capacity.

CATIP-engineered tumor-cell vaccination extends this principle. By covalently decorating tumor cells before implantation, CATIPs converted poorly immunogenic tumor cells into immunostimulatory whole-cell vaccines that prevented tumor establishment and instead induced durable tumor-specific protection. This approach parallels emerging efforts to transcriptionally reprogram tumor cells into APC-like cells,^29–31^ but differs in both mechanism and implementation. Genetic reprogramming strategies require efficient delivery and sustained expression of transcriptional regulators to induce new cell states, processes that can be heterogeneous and time-dependent. By contrast, covalent surface engineering installs defined immunostimulatory ligands and cytokines directly at the tumor-cell surface. This protein-level modification preserves endogenous tumor antigens and enables immediate functional activation without altering tumor-cell identity. Covalent surface engineering may therefore provide a modular alternative to genetic reprogramming for generating immunostimulatory whole-cell tumor vaccines.

The CATIP platform is not limited to the combinations used for efficacy studies. The broader CATIP library indicates that covalent tumor anchoring can be extended to multiple immunomodulatory pathways, including checkpoint blockade, innate immune activation and myeloid-directed signaling. This modularity is important because effective solid-tumor immunity rarely depends on a single immune axis. CATIPs therefore provide a means to assemble defined combinations of immune signals directly at tumor-cell surfaces, rather than distributing each component systemically.

This tumor-surface engineering strategy also differs from cell-intrinsic immune-cell engineering approaches. CAR T cells and related approaches endow transferred lymphocytes with tumor-recognition and co-stimulatory functions, but their activity is primarily mediated by transferred cells that must infiltrate tumors and function within immunosuppressive microenvironments.^5,69^ CATIPs instead reprogram the tumor-cell surface in situ, enabling anchored ligands to engage endogenous or transferred immune cells already present in the tumor microenvironment. This distinction may allow local immune activation to extend beyond a predefined engineered cell population and involve multiple immune compartments.^5,70,71^

Several limitations should be considered. The efficacy studies used EGFR-high tumor models, which likely favored anchoring and retention; whether comparable activity can be achieved with lower-density or heterogeneous targets will require further evaluation. Consistent with this, the exclusion of HER2 due to rapid internalization highlights the importance of target selection, as receptor abundance, turnover and trafficking are likely to influence both anchoring efficiency and therapeutic outcome. The cellular mechanisms underlying antigen spreading, abscopal tumor control and memory formation also remain to be defined in greater detail.^71^ Humanized NSG models capture key aspects of human immune activation but do not fully recapitulate its complexity. Finally, although intratumoral delivery is clinically feasible,^13^ future studies should determine how tumor size, anatomical site, target density and dosing influence biodistribution and therapeutic outcome.

In summary, CATIPs define a framework for engineering immune signaling at tumor-cell surfaces by controlling where immunostimulatory proteins reside and how long they are retained. Rather than increasing systemic exposure to intensify immune activation, covalent tumor anchoring increases the effective dwell time and organization of immune cues at tumor-cell interfaces. This enables local tumor eradication, antigen-specific systemic immunity, and durable protective memory while limiting systemic immunopathology. More broadly, these findings establish covalent tumor anchoring as a general strategy for spatially orchestrating antitumor immunity with enhanced precision, potency and safety.

## ACKNOWLEDGEMENTS

L.W. acknowledges support from the National Institutes of Health (NIH) (R01GM118384 and R01CA258300).

## Supplementary Information

Experimental materials and methods; Table S1-S4; Figure S1-S10.

