## Supplementary Information for "Covalent tumor anchoring spatially orchestrates antitumor immunity"

### EXPERIMENTAL MODEL AND DETAILS

#### Mice

For all experiments, female nude mice (F*oxn1*$^{nu}$; Strain #:002019), NSG mice (NOD.Cg-*Prkdc*$^{scid}$ *Il2rg*$^{tm1Wjl}$/SzJ; Strain #:005557) and C57BL/6J mice (Strain #000664), aged 6–8 weeks at the start of the initial experiment, were purchased from Jackson Laboratory and housed in the CVRI Animal Facility at UCSF. All animal procedures were performed in accordance with a protocol approved by the UCSF Institutional Animal Care and Use Committee (IACUC; protocol #AN202773-00E).

#### Cell lines

HEK-293T, MDA-MB-231 (human triple-negative breast cancer cells), A431 (human epidermoid carcinoma cells) and MDA-MB-468 (human breast cancer cells) were purchased from ATCC. Murine B16F10 melanoma cells, B16F10-OVA cells and murine colon adenocarcinoma MC38 cells were obtained from Eyquem lab (UCSF). B16F10-OVA-hEGFR/PD-L1 cells were generated by our lab. All these cells were cultured in DMEM (FISHER SCIENTIFIC, 11965092) supplemented with 10% FBS (GIBCO, 10099141) and penicillin-streptomycin (100U/ml) (DMEM complete medium). Human PBMCs were purchased from Charles River Laboratories (PB009C-2) and cultured in RPMI 1640 (FISHER SCIENTIFIC, 11875093) supplemented with 10% FBS and penicillin-streptomycin (100U/ml) (RPIM 1640 complete medium).

### METHOD DETAILS

#### Cloning of Tumor Immunotherapeutic Proteins (TIPs)

All synthetic DNA fragments encoding TIPs were cloned into the pET-22b vector with a C-terminal 6×His tag and Flag tag. Briefly, codon-optimized gene fragments (Integrated DNA Technologies, IDT) were ligated into NcoI- and XhoI-digested pET-22b vectors using the ClonExpress® Ultra One Step Cloning Kit (Vazyme, C115-02).

To introduce TAG codons at specific positions, site-directed mutagenesis was performed. The TAG codon was introduced at Y109 of $Nb_{EGFR}$, D54 of $Nb_{HER2}$.

Amino acid sequences of all TIPs used in this study are listed in **Table 1**.

#### Expression and purification of TIPs

Plasmids pET-22b-TIPs (WT) were transformed into E. coli BL21(DE3) competent cells. Plasmid pET-22b-$Nb_{EGFR}$(Y109TAG)-TIPs or pET-22b-$Nb_{HER2}$(D54TAG)-TIPs was co-transformed with plasmid pEvol-FSYRS (Wang et al., 2018) into E. coli BL21(DE3) competent cells. For expression of WT-TIPs, transformed cells were cultured at 37 °C in Super Broth medium (32 g tryptone, 20 g yeast extract, and 5 g NaCl per liter of $H_2O$) supplemented with 100 µg/mL ampicillin. When the culture reached an $OD_{600}$ of approximately 0.8, protein expression was induced with 1 mM IPTG, and incubation was maintained at 18–22 °C for protein expression. For expression of CATIPs, transformed cells were cultured at 37 °C in Super Broth medium supplemented with 100 µg/mL ampicillin and 34 µg/mL chloramphenicol. When the culture reached an $OD_{600}$ of approximately 0.8, CATIPs expression was induced by adding 1 mM IPTG, 0.2% (w/v) L-arabinose, and 1 mM FSY. The cultures were then incubated at 18–22 °C for protein expression. After induction of expression for 18-22 h, cells were collected by centrifugation at 8000 rpm for 5 min at 4 °C. The cell pellets were re-suspended in 10 mL lysis buffer (20 mM Tris-HCl, pH 8.0, 200 mM NaCl, 0.5% Triton X-100, lysozyme 1 mg/mL, DNase 0.1mg/mL, and protease inhibitors) per gram of bacteria wet weight. The cell suspension was lysed at 4 °C for 30 min by shaking and subsequently sonicated in an ice-water bath. The supernatant was then recovered by centrifugation of the cell lysate at 20,000 g for 20 min at 4 °C. Appropriate volume of Ni-NTA Resin (Thermo Scientific, 88221) were added into the supernatant and incubated at 4 °C for 0.5-1 hour by shaking. The mixture was loaded in a column and washed with wash buffer (20 mM TrisHCl, pH 8.0, 400 mM NaCl, 20 mM imidazole) in 10 column volume. The protein was eluted with elution buffer (20 mM TrisHCl, pH 8.0, 400 mM NaCl, 300 mM imidazole). The eluted protein solution was loaded into an ultrafiltration centrifugation tube (Millipore, UFC501024, MWCO = 10 kDa) to concentrate and exchange buffer into PBS (pH 7.4) (for in vitro protein crosslinking, cell and animal experiments). The concentrated and buffer-exchanged TIPs were stored at −80 °C and sterilized by filtration through a 0.22 μm filter prior to use in cell- and mouse-based experiments. The purity of all TIPs used for in vivo experiments exceeded 95%. Protein concentration was measured by nanodrop based on its molecular weight and extinction coefficient. The predicted molecular weight and extinction coefficient of all TIPs are listed in **Table 1**.

**Crosslinking of CATIPs with EGFR ECD in vitro**

Purified $Nb_{EGFR}$-$Nb_{CD3}$, $Nb_{EGFR}$-CD80, or $Nb_{EGFR}$-Neo2/15 was incubated with EGFR ECD at the molar ratio of 1:1 in 10 μL PBS buffer at 37 °C for 16 h. The amount of EGFR ECD (Sino Biological, 10001-H08H) used was 6 μg (MW=110 kDa).

Purified $Nb_{EGFR}$(FSY)-$Nb_{CD3}$ was incubated with EGFR ECD at the molar ratio of 1:4 or 1:8 in 10 μL PBS buffer at 37 °C for 12 h. The amount of EGFR used was 2 or 4 μg, respectively.

Purified $Nb_{EGFR}$-21h10 was incubated with EGFR ECD at the molar ratio of 1:1 in 10 μL PBS buffer at 37 °C for 14h. The amount of EGFR ECD used was 6 μg.

After incubation, 4x reduced loading buffer (Bio-Rad, 1610747, 20 mM DTT) was added into the mixture and heated at 95 °C for 10 min. These samples were then separated by 10% SDS-PAGE gel followed by staining with Coomassie brilliant blue. Western blot was then performed using the anti-6x His (Proteintech, HRP-66005) or anti-Flag (Proteintech, HRP-66008). Protein bands were detected by chemiluminescence (Thermo Scientific, 34577).

**Crosslinking of TIPs with TAA targets on cell surface**

A-431, MDA-MB-468, B16-F10-OVA-hEGFR/PD-L1 or B16-F10-OVA cells ($5x10^5$) were seeded in 12-well plates and cultured with complete DMEM medium, respectively. When cell confluence reached 70–80%, TIPs were added into wells in a final volume of 1 mL. After incubation at 37 °C for 12 h, cells were dissociated with 0.25% trypsin-EDTA, collected, and lysed by adding 100 μL of RIPA buffer (Thermo Scientific, 89900) with protease inhibitors (Roche, 11836170001) and DNase I (Roche, 10104159001), followed by incubation at 4 °C for 30 min. Protein concentrations were determined using a BCA protein assay kit (Thermo Scientific, 23227). Samples were then mixed with 4× reducing loading buffer (Bio-Rad, 1610747, 20 mM DTT) and heated at 70 °C for 10 min to denature the proteins. The denatured samples were separated on 7% SDS-PAGE gels and transferred for western blot analysis. Primary antibodies against human EGFR (Proteintech, 66455-1-Ig) or HER2 (CST, 4290S) was used, followed by an HRP-conjugated anti-mouse IgG (CST, 7076S) or anti-rabbit IgG (CST, 7074S). GAPDH (Proteintech, HRP-60004) was used as an internal control. Protein bands were visualized using chemiluminescence.

**CATIP retention and distribution in cells**

For in vitro crosslinking and consumption analysis at varying CATIPs concentrations, 5 × $10^5$ A-431 cells were seeded into each well of a 12-well plate and cultured to ~90% confluence. The medium was then replaced with 250 µL of fresh medium. $Nb_{EGFR}$-$Nb_{CD3}$ was added to final concentrations of 10 or 50 nM, and the total volume was adjusted to 300 µL. At the indicated time points, the supernatant was collected. Cells were washed twice with 100 µL PBS, and the wash fractions were combined with the corresponding supernatant. The combined solution was then adjusted to a final volume of 500 µL. Samples were mixed with denaturing loading buffer and heated at 95 °C for 10 min, followed by separation on 10% SDS-PAGE and subsequent western blot analysis using an HRP-anti-Flag antibody.

For in vitro crosslinking and consumption analysis at varying tumor cell numbers, 0.5 × $10^6$, 2.5 × $10^6$, or 10 × $10^6$ A-431 cells were seeded into each well of a 12-well plate. $Nb_{EGFR}$-$Nb_{CD3}$ was added to a final concentration of 25 nM, and the total reaction volume was adjusted to 300 µL. At the indicated time points, the supernatant was collected and the cells were washed twice with PBS. The harvested supernatant and wash fractions were then processed for western blot analysis as described above.

For analysis of the crosslinking of $Nb_{EGFR}$-CD80, 5 × $10^5$ A-431 cells were seeded into each well of a 12-well plate. After 6 h of cell adhesion, $Nb_{EGFR}$-CD80 was added to the cells at a final concentration of 100 nM (for time-dependent assays at a fixed concentration) or at the indicated concentrations (for dose–dependent assays at a fixed time point). After incubation at the specified time points, cells were washed three times with PBS, detached using 0.25% trypsin–EDTA, collected, and stained with PE-anti-Flag antibody (Biolegend, 637310) for 30 min at room temperature. After two additional PBS washes, cells were analyzed by flow cytometry.

For analysis of the in vitro crosslinking of $Nb_{EGFR}$-Neo2/15, 5 × $10^5$ A-431 cells were seeded into each well of a 12-well plate. When cell confluency reached approximately 90%, $Nb_{EGFR}$-Neo2/15 was added at a final concentration of 10 nM or 30 nM, and the total reaction volume was adjusted to 300 µL. After 6.5 h of incubation, the cells were washed three times with 100 µL PBS and then dissociated with 0.25% trypsin–EDTA, collected and stained with PE-anti-Flag antibody (Biolegend, 637310) for 30 min at 4 °C. The cells were washed twice with PBS and analyzed by flow cytometry. Flow cytometry data were analyzed using FlowJo v10.

**CATIP retention and distribution in tumor in mice**

A431 cells (10x$10^6$ for the primary tumor or 4x$10^6$ for the adjacent tumor) were resuspended with 100 µL PBS and injected into the flanks of 6-8 weeks old nude mice. After 18 days, the tumor volumes reached approximately 150 $mm^3$ for the primary tumors and 50 $mm^3$ for the adjacent tumors. Either high-dose (15 µg per primary tumor at the right flank) or low-dose (1.5 µg per primary tumor at the left flank) of $Nb_{EGFR}$(FSY)/(WT)-$Nb_{CD3}$ was intratumorally administered into the primary A431 tumors. A total of four doses were administered at 12-hour intervals. Eight hours after the final dose, blood was collected into heparin-containing tubes for plasma isolation, and tumors were harvested. The plasma samples were then stored at –80 °C. These harvested tumor tissues were cut into 1-2 mm patches and dissociated by adding 3 mL tissue digestive enzyme solution [100 Kunitz units of DNaseI (Roche, 10104159001), Collagenase D (10 U/mL, Roche, 11088866001) in FBS-free RPMI-1640 medium]. After shaking at 37 °C for 30 min, the digestion was stopped by adding 10 mL cold RPMI-1640 medium containing 10% FBS. Subsequently, the resulting cell suspension was filtered through 40 μm filter and spun down. The supernatant was aspirated, and the cells were resuspended in PBS and divided into two portions. One portion was stained with PE-anti-Flag antibody for 30 min and then analyzed by flow cytometry. The other portion was lysed by adding 200 μL RIPA buffer with protease inhibitors and DNase I. Lysates were processed for western blotting using the same procedures as described in the section above for detection of EGFR crosslinking.

Collected plasma were serially diluted 0-, 3-, 9-, 27-, 45-, 63-, 81-, 99-, and 117-fold with PBS, and then incubated with fresh A431 cells at 4 °C for 30 minutes. After incubation, the cells were washed once with PBS, stained with PE–anti-Flag antibody for 30 minutes at 4 °C. Cells were then washed again with PBS and analyzed by flow cytometry. Flow cytometry data were analyzed using FlowJo v10.

**Evaluation of TIPs activity in vitro**

To evaluate the T cell activation and PBMC-toxicity enhanced by $Nb_{EGFR}$-$Nb_{CD3}$, 1 × $10^4$ A431 or MDA-MB-468 cells in 100 μL complete DMEM medium were seeded per well in a 96-well plate. After 6 hours of cell adhesion, $Nb_{EGFR}$-$Nb_{CD3}$ was added at the indicated concentrations. Following a 12-hour incubation, the medium was either removed (PBS-washed group) or retained (non-washed group). For the PBS-washed group, cells were washed three times with 200 μL PBS. Subsequently, 6 × $10^4$ PBMCs in 100 μL complete RPMI-1640 medium were added to both the PBS-washed and non-washed groups. Co-culture was maintained for 72 hours for the PBS-washed group and 48 hours for the non-washed group.

After co-incubation, the suspended cells were collected for flow cytometry analysis of T cell (V450-anti-CD3 antibody, BD, 560366) activation by detecting the upregulation of CD25 (APC-anti-CD25, BD, 567316), CD69 (PE-Cy7-anti-CD69, BD, 561928), PD-1 (FITC-anti-PD-1, BD, 557860), and 4-1BB (PE-anti-4-1BB, BD, 561701). Flow cytometry data were analyzed using FlowJo v10.

For assessment of tumor cell viability, adherent A431 or MDA-MB-468 cells after co-culture were washed three times with PBS, followed by addition of 100 μL complete DMEM and 10 μL CCK-8 reagent (GLPBIO, GK10001) per well. After 1 hour of incubation at 37°C, cell viability was measured. The percentage of tumor cell viability was calculated as: viability (%) = {[OD450 of (tumor cells + PBMCs + TIPs) group -OD450 of medium only group)]/ [OD450 of (tumor cells + PBMCs + PBS) group]-OD450 of medium only group)} x 100.

To assess the activity of $Nb_{EGFR}$-CD80, PD-L1 expression on A431 and MDA-MB-468 cells was assessed, as CD80-mediated CD28 activation depends on CD80–PD-L1 interactions. Cells (5 × $10^5$) were seeded per well in a 6-well plate. When cells reached approximately 50% confluence, human IFN-γ (Sino Biological, 11725-HNAS) was added at final concentrations of 5, 50, or 100 ng/mL to induce upregulation of PD-L1 on the tumor cell surface. Subsequently, cell lysates were prepared and processed for western blotting following the procedures described above. Proteins were detected using the primary anti-human PD-L1 (Abcam, ab213524) and the second anti-rabbit IgG, with GAPDH serving as the internal loading control.

To assess T cell activation and PBMC-cytotoxicity enhanced by $Nb_{EGFR}$-CD80 or $Nb_{EGFR}$-Neo2/15 alone or in combination with $Nb_{EGFR}$-$Nb_{CD3}$, 1 × $10^4$ A431 cells in 100 μL complete DMEM were seeded per well in a 96-well plate. After 6 hours of cell adhesion, $Nb_{EGFR}$-$Nb_{CD3}$ was pre-added at the indicated concentrations 3 hours prior to the addition of $Nb_{EGFR}$-CD80 or $Nb_{EGFR}$-Neo2/15 in combination groups. This pre-incubation was performed to prevent complete EGFR occupancy by the higher concentration of $Nb_{EGFR}$-CD80 or $Nb_{EGFR}$-Neo2/15 relative to lower concentration of $Nb_{EGFR}$-$Nb_{CD3}$. After the three hours pre-incubation, $Nb_{EGFR}$-CD80 or Nb-Neo2/15 was added to the appropriate wells. Following a 12-hour incubation, the medium was either removed (PBS-washed condition) or retained (non-washed condition) and then $6 \times 10^4$ PBMCs were added using the same procedures as described for $Nb_{EGFR}$-$Nb_{CD3}$ alone. Co-cultures were maintained for 72 hours for the PBS-washed conditions and 60 hours for the non-washed conditions before flow cytometric and cell viability analyses.

**Human PBMC-reconstituted NSG mouse xenograft model study**

On day -24, 10 x $10^6$ A-431 cells mixed with 1x$10^6$ partially HLA-matched PBMCs and 10 x$10^6$ MDA-MB-468 cells mixed with 0.5 x $10^6$ partially HLA-matched PBMCs were separately injected into adjacent sites on the right flanks of 6–8-week-old NSG mice, generating two neighboring tumors per mouse. Co-injection of tumor cells with their respective partially HLA-matched PBMCs was intended to mitigate xenogeneic graft-versus-host disease (xGVHD)–associated nonspecific tumor killing during early tumor establishment. By day 0, A-431 tumors had grown to approximately 150 $mm^3$, while MDA-MB-468 tumors reached ~50 $mm^3$. Human PBMCs ($3 \times 10^6$) partially HLA-matched to MDA-MB-468 cells were adoptively transferred into tumor-bearing mice via tail vein injection. Concurrently, low- or high-dose TIPs combinations of WT or FSY variants—$Nb_{EGFR}$(WT)-$Nb_{CD3}$ + $Nb_{EGFR}$(WT)-CD80 + $Nb_{EGFR}$(WT)-Neo2/15 or $Nb_{EGFR}$(FSY)-$Nb_{CD3}$ + $Nb_{EGFR}$(FSY)-CD80 + $Nb_{EGFR}$(FSY)-Neo2/15—were administered intratumorally into A-431 tumors every three days.

For the low-dose group: $Nb_{EGFR}$-$Nb_{CD3}$ = 25 nM, $Nb_{EGFR}$-CD80 = 100 nM, and $Nb_{EGFR}$-Neo2/15 = 50 nM. For the high-dose group: $Nb_{EGFR}$-$Nb_{CD3}$ = 250 nM, $Nb_{EGFR}$-CD80 = 1000 nM, and $Nb_{EGFR}$-Neo2/15 = 500 nM. For example, a tumor with a volume of 200 $mm^3$ would receive 0.16 μg $Nb_{EGFR}$-$Nb_{CD3}$, 0.58 μg $Nb_{EGFR}$-CD80, and 0.29 μg $Nb_{EGFR}$-Neo2/15, for a total of 1.03 μg TIPs in the low-dose combination.

As a positive control, recombinant proteins comprising $Nb_{EGFR}$-$ScFv_{UCHT1}$, CD80, and Neo2/15—at molar concentrations equivalent to those used in the high-dose TIPs—were also injected intratumorally into A-431 tumors every three days.

Tumor growth was monitored using calipers, and tumor volume was calculated using the formula tumor volume = length × $width^2$ / 2. Relative changes of body weight were calculated using the formula (body weight at day x / body weight at day 0) x 100%. xGVHD was assessed using a modified clinical scoring system (see **Table 2**).

On day 33, all surviving mice were euthanized for downstream analyses. Tumors were excised and weighed. Peripheral blood was collected into heparinized tubes, and plasma was isolated and stored at −80 °C for human cytokine quantification by ELISA. Red blood cells were lysed twice using ACK lysis buffer (Quality Biological, 118-156-101), and the remaining cells were stained with anti-human CD3, CD8 (APC-anti-CD8, BD, 566852), CD69, PD-1, and 4-1BB antibodies for flow cytometry analysis. Flow cytometry data were analyzed using FlowJo v10.

Spleens were harvested and mechanically dissociated using syringe plungers. The resulting cell suspensions were resuspended in PBS, filtered through a 40 μm cell strainer (Corning, Cat# 352340), and centrifuged. Red blood cells were then lysed once using ACK lysis buffer to obtain splenocytes, which were subsequently stained and analyzed as described for blood samples.

Plasma supernatants were diluted as appropriate fold and analyzed using ELISA kits specific for human IFN-γ (Biolegend, 430104), TNF-α (Biolegend, 430204), IL-2 (431804), IL-6 (Biolegend, 430504), IL-10 (Biolegend, 430604), and GM-CSF (Biolegend, 432004), according to the manufacturers' instructions.

Collected liver tissues were subjected to H&E staining and histopathological imaging to assess GVHD severity and off-tumor toxicity–associated tissue damage.

Peripheral blood samples and spleens from deceased mice were also collected for flow cytometric and ELISA analyses.

**Immunocompetent mice to assess therapeutic efficacy, abscopal tumor regression, tumor-antigen specificity, and immune memory**

**Lentivirus preparation to generate B16-F10-OVA-hEGFR/PD-L1 cell line**

To construct the core plasmid encoding human EGFR and PD-L1, a codon-optimized DNA fragment, Kozak–mouse EGFR signal peptide (SP)–HA tag/human EGFR–IRES–mouse PD-L1 SP–Flag tag/human PD-L1, was synthesized by IDT and subsequently ligated into the pLenti-CMV-luciferase-SV40-BSD backbone (Addgene, 21474) digested by BamHI and XbaI. The amino acid sequences of overexpressed human EGFR and PD-L1 are listed in **Table 3**.

To prepare lenti-virus for transducing B16F10-OVA cell, HEK-293T cells were seeded in 10 cm cell dish cultured with DMEM containing 10% FBS (without antibiotics) at the confluency of ~50%. About 24 h later, cells grew to ~80% confluency. Ten μg core plasmid, 10 μg pMD2.G, and 15 μg psPAX2 were added into 1 mL Opti-MEM (GIBCO, Cat# 31985088), followed with addition of 105 μg polyetherimide (Sigma, Cat# GF70215825) (1 mg/mL). The mixture was vortexed for 20 s and protected from light for 15 min. The mixture was then added into HEK-293FT cell dish and mixed well. After 10 h, cell medium was replaced with fresh DMEM containing 10% FBS (without antibiotics). After 48 h, supernatant containing lenti-virus was harvested and centrifuged at 1500 rpm for 10 min to remove cell debris. The supernatant was prepared for transduction.

**Transduction of B16F10-OVA cells and confirmation of human EGFR and PD-L1 expression**

B16F10-OVA cells were seeded into a 6-well plate and 8 μg/mL polybrene (EMO Millipore, Cat# TR-1003-G) was added to enhance viral transduction. Lentivirus supernatant was added into the wells. After 12 h of infection, the medium was replaced with fresh complete DMEM. A second transduction was performed after 48 h. Transduced cells were selected with BSD (Sigma, SBR00022-1ML) at 3–10 μg/mL. After 10 days of selection, surviving cells were maintained in 7 μg/mL BSD. Selected cells were subsequently analyzed by western blot using anti-human EGFR and anti-PD-L1 antibodies. Additionally, flow cytometry was performed to confirm PD-L1 expression on the cell surface using PE-anti-Flag antibody.

**Co-culture of mouse splenocytes with B16F10-OVA-hEGFR/PD-L1 cells**

To evaluate the mouse splenocyte-cytotoxicity enhanced by $Nb_{EGFR}$-CD80, $Nb_{EGFR}$-Neo2/15 and

$Nb_{EGFR}$-21h10, 0.5 x $10^4$ B16F10-OVA-hEGFR/PD-L1 in 100 μL complete DMEM medium were seeded per well in a 96-well plate. After 6 hours of cell adhesion, TIPs were added at the indicated concentrations. Following a 12-hour incubation, the medium was either removed (PBS-washed) or retained (non-washed).

For the PBS-washed condition, cells were washed three times with PBS. Subsequently, 5 × $10^4$ mouse splenocytes in 100 μL complete RPMI-1640 medium were added to both the PBS-washed and non-washed groups. Co-culture was maintained for 72 hours for the PBS-washed condition and 48 hours for the non-washed condition. After co-incubation, adherent B16F10-OVA-hEGFR/PD-L1 cells were washed three times with PBS, followed by addition of 100 μL complete DMEM and 10 μL CCK-8 reagent per well. After 1 hour of incubation at 37 °C, cell viability was measured. The percentage of cell viability was calculated as: viability (%) = {[OD450 of (tumor cells + splenocytes + TIPs) group -OD450 of medium only group)]/ [OD450 of (tumor cells + splenocytes + PBS) group]-OD450 of medium only group)} x 100.

**In vivo evaluation of TIPs therapeutic efficacy in primary tumors and abscopal effect model**

On day -9, a total of 0.5 × $10^6$ B16F10-OVA-hEGFR/PD-L1 cells, 0.2 × $10^6$ B16-F10-OVA cells, and 1 × $10^6$ MC38 cells in 100 μL PBS were separately injected into the flanks of 6–8-week-old C57BL/6J mice, generating three tumors per mouse. Tumors were designated as the primary tumor (right posterior flank), distant untreated tumor (left posterior flank), and antigen-unrelated tumor (left anterior flank), respectively. This model was used to assess local therapeutic efficacy, the abscopal effect, and immunotherapy specificity or off-treated tumor toxicity. On day 0, when primary tumors reached ~150 mm³, low- or high- dose of WT-TIPs or CATIP combinations were administered intratumorally every three days.

For the low-dose group: $Nb_{EGFR}$-CD80 = 100 nM, $Nb_{EGFR}$-Neo2/15 = 100 nM, and $Nb_{EGFR}$-21h10 = 100 nM. For the high-dose group: $Nb_{EGFR}$-CD80 = 1000 nM, $Nb_{EGFR}$Neo-2/15 = 1000 nM, and $Nb_{EGFR}$-21h10 = 1000 nM. As an example, a tumor with a volume of 200 mm³ would receive 0.58 μg $Nb_{EGFR}$-CD80, 0.58 μg $Nb_{EGFR}$- Neo2/15, and 0.58 μg $Nb_{EGFR}$-21h10, for a total of 1.74 μg in the low-dose combination.

Tumor growth in all three sites per mouse was measured in two dimensions using calipers, and tumor volume was calculated as: volume = length × width² / 2.

Mice that developed large ulcerated tumors were euthanized. On day 12, surviving mice were euthanized for downstream analyses. Tumors were excised and weighed. Harvested tumors were then mechanically dissociated using syringe plungers, re-suspended in PBS and filtered through 40 μm filter and divided into two portions. One portion was stained with anti-mouse CD45 (PerCP/Cy5.5, Biolegend, 103131), CD3 (BV421, Biolegend, 100335), CD8a (APC, Biolegend, 100711), and PE-labeled mouse H-2Kb&B2M&OVA (SIINFEKL) tetramer protein (ACRO, H2A-MP2H7) for flow cytometry analysis, while the other was stored at –80 °C for RNA extraction.

Peripheral blood was collected into heparinized tubes. Red blood cells were lysed twice using ACK lysis buffer, and the remaining leukocytes were stained as the same with tumor samples for flow cytometry analysis. Data were analyzed using FlowJo v10.

For RNA analysis, primary tumor cells were lysed in 1 mL Trizol (Invitrogen, AM9738) and total RNA extracted using an RNA extraction kit (ZYMO research, R2050). Total RNA (1 μg) of each sample was reverse transcribed using first-strand cDNA synthesis kit (APExBio, K1072). The resulting cDNA was diluted threefold with DNase- and RNase-free water, and 2 μL of diluted cDNA was used per qPCR reaction with SYBR Green (Vazyme, Q712-02) on a Real-Time PCR system (CFX Opus 96). Relative mRNA expression of each gene was calculated using the following equation: mRNA level of gene X relative to β-actin = -ΔΔCT, wherein $\Delta\Delta CT = (CT_{gene\ X,\ drug} - CT_{\beta\text{-actin},\ drug}) - (CT_{gene\ X,\ PBS} - CT_{\beta\text{-actin},\ PBS})$. -ΔΔCT were used for heatmap. Primers for qRT-PCR are listed in **Table 4.**

**Assessment of tumor-antigen specific, durable immune memory following subcutaneous rechallenge in tumor-eliminated survivors**

To assess whether multiple administrations of high-dose CATIPs could completely eradicate established tumors and induce off-tumor toxicity, mice bearing only primary tumors were intratumorally treated with high-dose CATIPs in a total of 6–10 doses. Briefly, on day -9, a total of $0.5 \times 10^6$ B16F10-OVA-hEGFR/PD-L1 cells in 100 µL PBS were injected into the right posterior flank of 6–8-week-old C57BL/6J mice, generating the primary tumors. On day 0, when primary tumors reached ~150 mm³, high-dose CATIPs were administered intratumorally every three days.

For high-dose CATIPs: $Nb_{EGFR}$(FSY)-CD80 = 1000 nM, $Nb_{EGFR}$(FSY)-Neo2/15 = 1000 nM, and $Nb_{EGFR}$(FSY)-21h10 = 1000 nM. As an example, a tumor with a volume of 200 mm³ would receive 5.8 µg $Nb_{EGFR}$(FSY)-CD80, 5.8 µg $Nb_{EGFR}$(FSY)-Neo2/15, and 5.8 µg $Nb_{EGFR}$(FSY)-21h10, for a total of 17.4 µg CATIPs.

Tumor growth and body weight were monitored.

On day 66, $2 \times 10^5$ B16F10-OVA cells, $2 \times 10^5$ B16F10 cells, and $1 \times 10^6$ MC38 cells in 100 µL PBS were separately injected into the flanks of tumor-eliminated survivor mice. This model was designated as **Antigen-specificity model.** Tumor growth was subsequently monitored.

In parallel, to determine whether tumor-eradicated survivors had established durable immune protection, $0.2 \times 10^6$ B16-F10-OVA cells in 100 µL PBS were injected into the right anterior flank of the survivors. This model was designated as **Immune memory (s.c. rechallenge).** Naïve C57BL/6J mice were included as controls. Long-term survival was monitored.

**Evaluation of long-term survival following intravenous rechallenge in s.c. rechallenge-resistant survivors**

To determine whether tumor-free survivors post s.c. rechallenge had established durable immune protection against intravenous rechallenge and subsequent lung metastasis, on day 126, $0.2 \times 10^6$ B16F10-OVA cells in 100 µL PBS were injected into the tail vein of survivors from **Immune memory (s.c. rechallenge)**. This model was designated as **Durable immune memory against lung metastases (i.v. rechallenge)**. Naive C57BL/6J mice were included as controls. Long-term survival was monitored.

At the indicated time points upon death, lungs were harvested and fixed in 4% paraformaldehyde for 48 h, followed by imaging to assess B16-F10-OVA metastatic nodules.

On day 186, all surviving mice were photographed to document vitiligo-like depigmentation and localized hair whitening, and then euthanized for analysis. Spleens were harvested and mechanically dissociated using syringe plungers. The resulting cell suspensions were resuspended in PBS, and splenic mononuclear cells were isolated using Ficoll-Paque (Cytiva, 17544602). One-third of splenic mononuclear cells were then stained with anti-mouse CD45, CD3, CD8, PE-labeled mouse H-2Kb&B2M&OVA (SIINFEKL) tetramer protein, CD44 (FITC-anti-CD44, Biolegend, 103005), and CD62L (PE-Cy7-anti-CD62L, Biolegend, 104417) antibodies for flow cytometry analysis. Two-thirds of the splenic mononuclear cells were cryopreserved for subsequent ELISpot assays. The heart, liver, spleen, lungs, kidney and pancreas were collected for H&E staining to evaluate off-tumor toxicity associated with the repeated intratumoral administration of TIPs at the initial therapies.

For the ELISpot assay, $5 \times 10^5$ splenic mononuclear cells in 100 μL of complete RPMI 1640 medium were seeded per well in a 96-well mouse IFN-γ ELISpot plate. 100 μL complete RPMI 1640 containing OVA peptides (OVA257–264, Genscript C744B49; or OVA323–339, Genscript C744B47) was added to each well, resulting in a final peptide concentration of 5 μg/mL. Cells were incubated for 48 hours for recall stimulation. Following incubation, the ELISApot plate was developed according to the manufacturer's instructions. Spots were then imaged and quantified using an ELISpot reader.

**CATIPs-engineered whole tumor cell vaccines**

To generate CATIP-engineered whole tumor cell vaccines, B16F10-OVA-hEGFR/PD-L1 cells were seeded in 10-cm dishes. When cells reached approximately 70% confluency, $Nb_{EGFR}$(FSY)-CD80, $Nb_{EGFR}$(FSY)-Neo2/15 and $Nb_{EGFR}$(FSY)-21h10 combinations were added to the dishes at a final concentration of 200 nM in 12 mL of complete DMEM for covalent crosslinking and pre-engineering (CATIPs-V). WT-TIPs with equal molar concentrations to CATIPs were included as controls. After 12 h of incubation, the cells were washed twice with PBS, dissociated with 0.25% trypsin-EDTA, and collected.

For vaccination, $0.2 \times 10^6$ pre-engineered vaccine cells were subcutaneously injected into the right posterior flank of 6–8-week-old C57BL/6J mice. Mice were immunized with two, three, or four doses of CATIPs-V every four days. Control mice were immunized with four doses of WT-TIPs-V. Tumor growth and survival were monitored.

**Assessment of antigen-specific and durable immune memory rejecting subcutaneous rechallenge in tumor-free survivors immunized with CATIPs-V**

On day 60, a total of $0.2 \times 10^6$ B16F10-OVA cells, $0.2 \times 10^6$ B16F10 cells, and $1 \times 10^6$ MC38 cells in 100 μL PBS were separately injected into the flanks of survivors which had been previously immunized with CATIPs-V. Tumors were designated as the rechallenged OVA-positive tumor (right anterior flank), rechallenged OVA-negative tumor (left posterior flank), and antigen-unrelated tumor (left anterior flank), respectively. This model was designated as **Antigen-specificity model.** Tumor growth was subsequently monitored.

In parallel, to determine whether these immunized tumor-free survivors had established durable immune protection, $0.2 \times 10^6$ B16F10-OVA cells in 100 μL PBS were injected into the right flank of the survivors. This model was designated as **Immune memory (s.c. rechallenge).** Naïve C57BL/6J mice were included as controls. Long-term survival was monitored.

**Evaluation of long-term survival following lung metastasis rechallenge in s.c. rechallenge-resistant survivors**

To determine whether survivors post s.c. rechallenge had established durable immune protection against intravenous rechallenge potentially generating subsequent lung metastasis, on day 120, $0.2 \times 10^6$ B16F10-OVA cells in 100 μL PBS were injected into the tail vein of survivors from **Immune memory (s.c. rechallenge)**. This model was designated as **Durable immune memory against lung metastases (i.v.**

**rechallenge)**. Naive C57BL/6J mice were included as controls. Long-term survival was monitored.

At the indicated time points upon death, lungs were harvested and fixed in 4% paraformaldehyde for 48 h, followed by imaging to assess B16-F10-OVA metastatic nodules.

On day 180, all surviving mice were photographed to document vitiligo-like depigmentation and localized hair whitening, and then euthanized for analysis. Spleen samples were collected and processed as described above. Splenic mononuclear cells were also used for ELISApot assays as described above.

**Table S1. Information for TIPs**

| Name | Amino acid sequence | Predicted molecular weight (Da) | Predicted extinction coefficient | Refs. |
|---|---|---|---|---|
| **Nb$_{EGFR}$**-Nb$_{CD3}$ | MGQVKLEESGGGSVQTGGSLRLTCAAS<br>GRTSRSYGMGWFRQAPGKRREFVSGIS<br>WRGDSTGYADSVKGRFTISRDNAKNTV<br>DLQMNSLKPEDTAIYYCAAAAGSAWY<br>GTLUEYDYWGQGTQVTVSSGGGGSGG<br>GGSGGGGSGGGGSEVQLVESGGGPVQ<br>AGGSLRLSCAASGRTYRGYSMGWFRQ<br>APGKEREFVAAIVWSGGNTYYEDSVKG<br>RFTISRDNAKNTMYLQMTSLKPEDSAT<br>YYCAAKIRPYIFKIAGQYDYWGQSTQV<br>TVSSHHHHHHDYKDDDDK.<br>U=Y for WT; U=FSY for CATIPs | 30290 | 67060 | 1 |
| Nb$_{EGFR}$-CD80 | MGQVKLEESGGGSVQTGGSLRLTCAAS<br>GRTSRSYGMGWFRQAPGKRREFVSGIS<br>WRGDSTGYADSVKGRFTISRDNAKNTV<br>DLQMNSLKPEDTAIYYCAAAAGSAWY<br>GTLUEYDYWGQGTQVTVSSGGGGSGG<br>GGSGGGGSGGGGSVIHVTKEVKEVATL<br>SCGYNVSVEELEQTRIYWQKDKKMVL<br>TMMSGDLNIWPEYKNRTIFDITNNLSIM<br>ILGLRPSDEGTYECVVLKYEKGAFKRE<br>HLAEVTLSVKAHHHHHHDYKDDDDK. | 28631 | 52620 | 2 |
| Nb$_{EGFR}$-Neo2/15 | MGQVKLEESGGGSVQTGGSLRLTCAAS<br>GRTSRSYGMGWFRQAPGKRREFVSGIS<br>WRGDSTGYADSVKGRFTISRDNAKNTV<br>DLQMNSLKPEDTAIYYCAAAAGSAWY<br>GTLUEYDYWGQGTQVTVSSGGGGSGG<br>GGSGGGGSGGGGSGSHMPKKKIQLHA<br>EHALYDALMILNIVKTNSPPAEEKLEDY<br>AFNFELILEEIARLFESGDQKDEAEKAK<br>RMKEWMKRIKTTASEDEQEEMANAIIT<br>ILQSWIFSHHHHHHDYKDDDDK. | 28661 | 49515 | 3 |
| Nb$_{EGFR}$-21h10 | MGQVKLEESGGGSVQTGGSLRLTCAAS<br>GRTSRSYGMGWFRQAPGKRREFVSGIS<br>WRGDSTGYADSVKGRFTISRDNAKNTV<br>DLQMNSLKPEDTAIYYCAAAAGSAWY | 28725 | 46535 | 4 |

| | | | | |
|---|---|---|---|---|
| | GTLUEYDYWGQGTQVTVSSGGGGSGGGGSGGGGSGGGGSDEEELKEIMKEAAECARKELEKLDNTDEDTRWLKITLKKIVRMANRIVRMREREADSKRFEIRMRGLIDIADHVKREFASEDLKEVMERAKSAAQKALGRWLHHHHHHDYKDDDDK. | | | |
| $Nb_{EGFR}$-$ScFv_{UCHT1}$ | MGQVKLEESGGGSVQTGGSLRLTCAASGRTSRSYGMGWFRQAPGKRREFVSGISWRGDSTGYADSVKGRFTISRDNAKNTVDLQMNSLKPEDTAIYYCAAAAGSAWYGTLYEYDYWGQGTQVTVSSGGGGSGGGGSGGGGSGGGGSMGMDIQMTQTTSSLSASLGDRVTISCRASQDIRNYLNWYQQKPDGTVKLLIYYTSRLHSGVPSKFSGSGSGTDYSLTISNLEQEDIATYFCQQGNTLPWTFAGGTKLEIKGGGGSGGGGSGGGGSEVQLQQSGPELVKPGASMKISCKASGYSFTGYTMNWVKQSHGKNLEWMGLINPYKGVSTYNQKFKDKATLTVDKSSSTAYMELLSLTSEDSAVYYCARSGYYGDSDWYFDVWGQGTTLTVFSHHHHHHDYKDDDDK. | 43304 | 92625 | 5 |
| CD80 | MGVIHVTKEVKEVATLSCGYNVSVEELEQTRIYWQKDKKMVLTMMSGDLNIWPEYKNRTIFDITNNLSIMILGLRPSDEGTYECVVLKYEKGAFKREHLAEVTLSVKAHHHHHHDYKDDDDK. | 14104 | 20065 | 2 |
| Neo2/15 | MGGSHMPKKKIQLHAEHALYDALMILNIVKTNSPPAEEKLEDYAFNFELILEEIARLFESGDQKDEAEKAKRMKEWMKRIKTTASEDEQEEMANAIITILQSWIFSHHHHHHDYKDDDDK. | 13970 | 15470 | 3 |

Protein sequences essential for interpretation of the main findings are provided here; additional supporting sequences have been made available for editorial review and will be publicly available upon publication.

**Table S2. Modified five-parameter clinical xGVHD scoring system.**

Each parameter is scored 0–2 points, with a total score of 0–10.

| **Parameter** | **0 (Normal)** | **1 (Moderate)** | **2 (Severe)** |
|---|---|---|---|
| Body weight change | < 5% loss | 5–10% loss | >10% loss |
| Posture | Normal | Mild hunching | Pronounced hunching |
| Activity | Normal | Mildly reduced | Markedly immobile |
| Fur condition | Smooth | Ruffled coat | Severely unkempt |
| Anemia | Normal coloration | Mild pallor (e.g., ears, paws, tail) | Marked pallor / severe anemia |

Total score interpretation:

0–2: No or minimal GVHD; 3–5: Mild GVHD; 6–8: Moderate GVHD; 9–10: Severe GVHD. Mice are euthanized when the clinical xGVHD score exceeds 8.

**Table S3. Amino acid sequences of human EGFR and PD-L1 expressed on B16F10-OVA-hEGFR/PD-L1 cells.**

| Protein | Amino acid sequence |
|---|---|
| Mouse EGFR signal peptide-**HA tag**-human EGFR | MRPSGTARTTLLVLLTALCAAGGA**YPYDVPDYA**LEEKKVCQGTSNKLTQ<br>LGTFEDHFLSLQRMFNNCEVVLGNLEITYVQRNYDLSFLKTIQEVAGYVLI<br>ALNTVERIPLENLQIIRGNMYYENSYALAVLSNYDANKTGLKELPMRNLQ<br>EILHGAVRFSNNPALCNVESIQWRDIVSSDFLSNMSMDFQNHLGSCQKCD<br>PSCPNGSCWGAGEENCQKLTKIICAQQCSGRCRGKSPSDCCHNQCAAGCT<br>GPRESDCLVCRKFRDEATCKDTCPPLMLYNPTTYQMDVNPEGKYSFGATC<br>VKKCPRNYVVTDHGSCVRACGADSYEMEEDGVRKCKKCEGPCRKVCN<br>GIGIGEFKDSLSINATNIKHFKNCTSISGDLHILPVAFRGDSFTHTPPLDPQEL<br>DILKTVKEITGFLLIQAWPENRTDLHAFENLEIIRGRTKQHGQFSLAVVSLN<br>ITSLGLRSLKEISDGDVIISGNKNLCYANTINWKKLFGTSGQKTKIISNRGE<br>NSCKATGQVCHALCSPEGCWGPEPRDCVSCRNVSRGRECVDKCNLLEGE<br>PREFVENSECIQCHPECLPQAMNITCTGRGPDNCIQCAHYIDGPHCVKTCP<br>AGVMGENNTLVWKYADAGHVCHLCHPNCTYGCTGPGLEGCPTNGPKIPS<br>IATGMVGALLLLLVVALGIGLFMRRRHIVRKRTLRRLLQERELVEPLTPSGE<br>APNQALLRILKETEFKKIKVLGSGAFGTVYKGLWIPEGEKVKIPVAIKELR<br>EATSPKANKEILDEAYVMASVDNPHVCRLLGICLTSTVQLITQLMPFGCLL<br>DYVREHKDNIGSQYLLNWCVQIAKGMNYLEDRRLVHRDLAARNVLVKT<br>PQHVKITDFGLAKLLGAEEKEYHAEGGKVPIKWMALESILHRIYTHQSDV<br>WSYGVTVWELMTFGSKPYDGIPASEISSILEKGERLPQPPICTIDVYMIMV<br>KCWMIDADSRPKFRELIIEFSKMARDPQRYLVIQGDERMHLPSPTDSNFYR<br>ALMDEEDMDDVVDADEYLIPQQGFFSSPSTSRTPLLSSLSATSNNSTVACI<br>DRNGLQSCPIKEDSFLQRYSSDPTGALTEDSIDDTFLPVPEYINQSVPKRPA<br>GSVQNPVYHNQPLNPAPSRDPHYQDPHSTAVGNPEYLNTVQPTCVNSTFD<br>SPAHWAQKGSHQISLDNPDYQQDFFPKEAKPNGIFKGSTAENAEYLRVAP<br>QSSEFIGA. |
| Mouse PD-L1 signal peptide-**Flag tag**-human PD-L1 | MRIFAGIIFTACCHLLRADYKDDDDKTYWHLLNAFTVTVPKDLYVVEYGS<br>NMTIECKFPVEKQLDLAALIVYWEMEDKNIIQFVHGEEDLKVQHSSYRQR<br>ARLLKDQLSLGNAALQITDVKLQDAGVYRCMISYGGADYKRITVKVNAP<br>YNKINQRILVVDPVTSEHELTCQAEGYPKAEVIWTSSDHQVLSGKTTTTNS<br>KREEKLFNVTSTLRINTTTNEIFYCTFRRLDPEENHTAELVIPELPLAHPPNE<br>RTHLVILGAILLCLGVALTFIFRLRKGRMMDVKKCGIQDTNSKKQSDTHLE<br>ET |

**Table S4. Primers for mouse genes used in qRT-PCR.**

| Gene name | Primer sequence |
|---|---|
| Actb | F: CATTGCTGACAGGATGCAGAAGG<br>R: TGCTGGAAGGTGGACAGTGAGG |
| Ifng | F: CAGCAACAGCAAGGCGAAAAAGG<br>R: TTTCCGCTTCCTGAGGCTGGAT |
| Il2 | F: GCGGCATGTTCTGGATTTGACTC<br>R: CCACCACAGTTGCTGACTCATC |
| Tnfa | F: GGTGCCTATGTCTCAGCCTCTT<br>R: GCCATAGAACTGATGAGAGGGAG |
| Csf2 | F: AACCTCCTGGATGACATGCCTG<br>R: AAATTGCCCCGTAGACCCTGCT |
| Il12a | F: ACGAGAGTTGCCTGGCTACTAG<br>R: CCTCATAGATGCTACCAAGGCAC |
| Gzmb | F: CAGGAGAAGACCCAGCAAGTCA<br>R: CTCACAGCTCTAGTCCTCTTGG |
| Prf1 | F: ACACAGTAGAGTGTCGCATGTAC<br>R: GTGGAGCTGTTAAAGTTGCGGG |
| Cxcl9 | F: CCTAGTGATAAGGAATGCACGATG<br>R: CTAGGCAGGTTTGATCTCCGTTC |
| Cxcl10 | F: ATCATCCCTGCGAGCCTATCCT<br>R: GACCTTTTTTGGCTAAACGCTTTC |
| Tnfrsf9 | F: CCAAGTACCTTCTCCAGCATAGG<br>R: GCGTTGTGGGTAGAGGAGCAAA |
| Cd69 | F: GGGCTGTGTTAATAGTGGTCCTC<br>R: CTTGCAGGTAGCAACATGGTGG |
| Cd80 | F: CCTCAAGTTTCCATGTCCAAGGC<br>R: GAGGAGAGTTGTAACGGCAAGG |
| Cd40 | F: ACCAGCAAGGATTGCGAGGCAT<br>R: GGATGACAGACGGTATCAGTGG |
| H2-K1 | F: GGCAATGAGCAGAGTTTCCGAG<br>R: CCACTTCACAGCCAGAGATCAC |
| H2-D1 | F: TGAGGAACCTGCTCGGCTACTA<br>R: GGTCTTCGTTCAGGGCGATGTA |
| B2m | F: ACAGTTCCACCCGCCTCACATT<br>R: TAGAAAGACCAGTCCTTGCTGAAG |
| Tap1 | F: GACTCCTTGCTCTCCACTCAGT<br>R: AACGCTGTCACCGTTCCAGGAT |
| Tap2 | F: AGCAGGAAGTCAGCCGCTACAA<br>R: CGCAGTTCAGAATCAGCACCTG |

| | |
|---|---|
| Tapbp | F: TGGTCAGCGTATCCAGCACTCT<br>R: TTATGGGTGAGGACGGTCAGCA |
| Foxp3 | F: CCTGGTTGTGAGAAGGTCTTCG<br>R: TGCTCCAGAGACTGCACCACTT |
| Tgfb1 | F: TGATACGCCTGAGTGGCTGTCT<br>R: CACAAGAGCAGTGAGCGCTGAA |

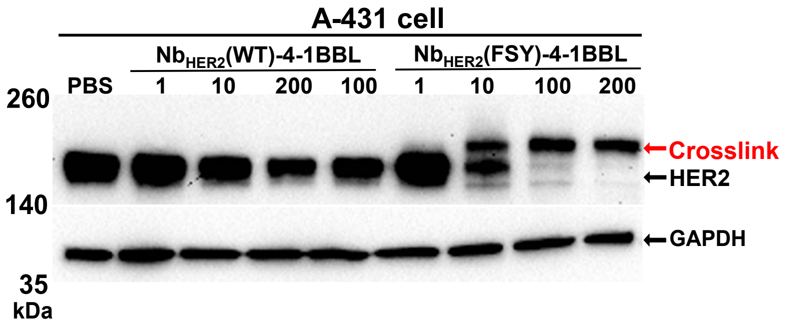


**Figure S1. HER2 targeted CATIPs induced HER2 degradation.**

Nb$_{HER2}$(FSY)-4-1BBL covalently bound HER2 on the surface of A431 cells, resulting in HER2 degradation in a dose-dependent manner.

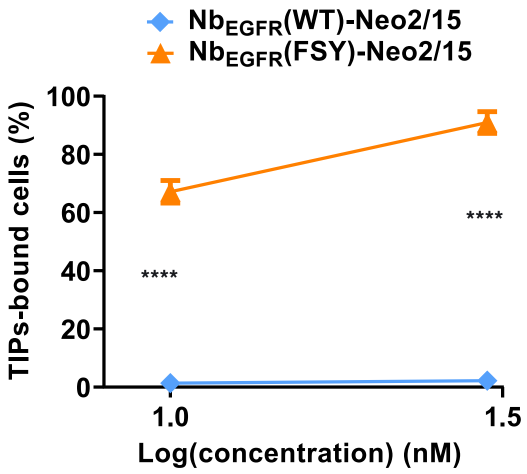


**Figure S2. $Nb_{EGFR}$(FSY)-Neo2/15 covalently bound A431 cells.**
$Nb_{EGFR}$(FSY)-Neo2/15 covalently bound EGFR on the surface of A431 cells in a dose-dependent manner. A431 cells were incubated with $Nb_{EGFR}$-Neo2/15 for 6.5 h, washed three times with PBS, and then detected for $Nb_{EGFR}$-Neo2/15 bound cells with flow cytometry. Data are mean ± SEM. n = 4 independent experiments. ****$P < 0.0001$; two-way ANOVA followed by Šídák's multiple-comparisons test.

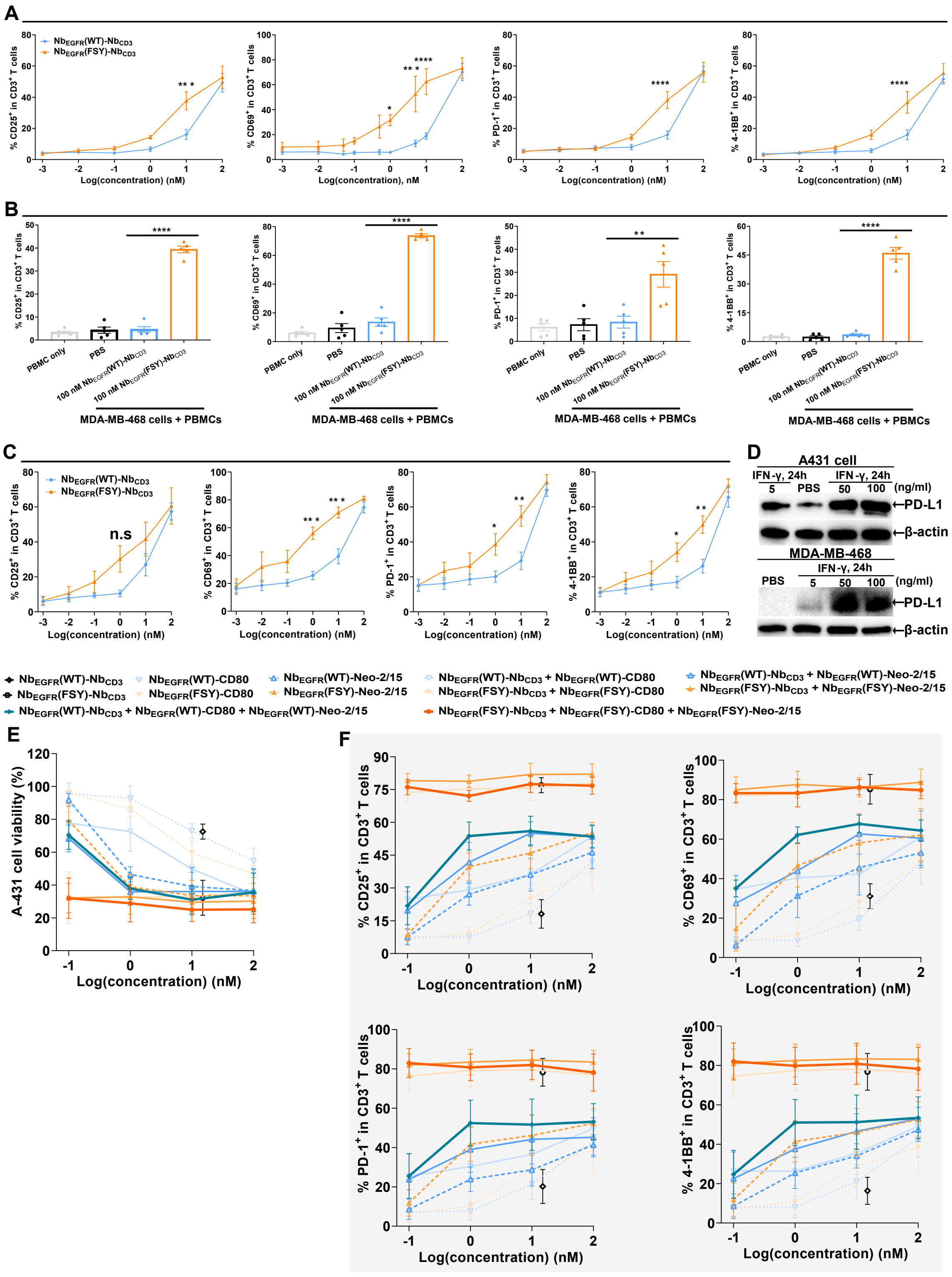

**Figure S3. CATIPs enhance T cell activation and PBMC-mediated cytotoxicity.**

(**A**) $Nb_{EGFR}$(FSY)–$Nb_{CD3}$ promoted T cell activation in a dose-dependent manner as shown by upregulation of the expression of indicated markers. A431 cells were pre-incubated with

$Nb_{EGFR}$(FSY)–$Nb_{CD3}$ under non-washed conditions and then co-cultured with PMBCs. Data are mean ± SEM. n = 4 independent experiments. **P* < 0.05; ****P* < 0.001; *****P* < 0.0001; two-way ANOVA followed by Šídák's multiple comparisons test.

(**B-C**) $Nb_{EGFR}$(FSY)–$Nb_{CD3}$ promoted T cell activation as shown by upregulation of the expression of indicated markers. MDA-MB-468 cells pre-incubated with $Nb_{EGFR}$(FSY)–$Nb_{CD3}$ under PBS-washed (B) or non-washed (C) conditions and then co-cultured with PBMCs. Data are mean ± SEM. n = 5 independent experiments. **P* < 0.05; ***P* < 0.01; ****P* < 0.001; *****P* < 0.0001; one-way ANOVA followed by Tukey's multiple comparisons test (B) or two-way ANOVA followed by Šídák's multiple comparisons test (C).

(**D**) Human IFN-γ induced PD-L1 expression on A431 and MDA-MB-468 cells in a dose-dependent manner.

(**E**) CATIP combinations cooperatively enhanced PBMC-mediated cytotoxicity against A431 cells under PBS non-washed conditions in a dose-dependent manner. The final concentration of $Nb_{EGFR}$–$Nb_{CD3}$ was 15 nM.

(**F**) CATIP combinations cooperatively promoted T cell activation as shown by upregulation of the expression of indicated markers. A431 cells were pre-incubated with CATIPs under PBS non-washed conditions and then co-cultured with PBMCs. The final concentration of $Nb_{EGFR}$–$Nb_{CD3}$ was 15 nM.

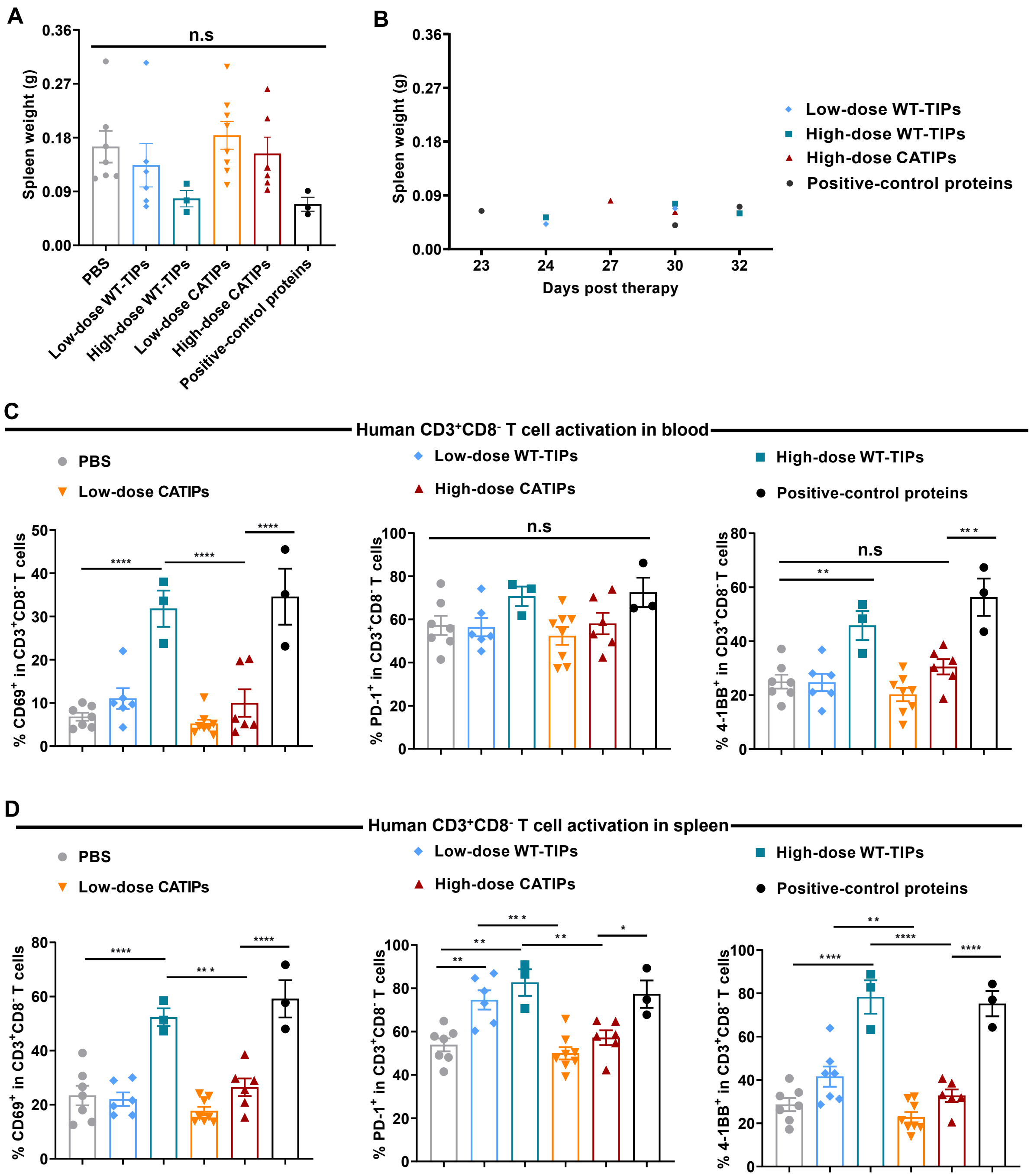


**Figure S4. CATIPs induced minimal human T cell activation.**

(**A-B**) Spleen weights at the study endpoint (A) and in deceased mice at the indicated time points (B). n.s. *P* > 0.05; one-way ANOVA followed by Tukey's multiple-comparisons test.

(**C-D**) Human $CD3^{+}CD8^{-}$ T cell activation in blood (C) or spleen (D) at the study endpoint. Data are mean ± SEM. n = 7 (PBS), 8 (Low-dose CATIPs), 6 (Low-dose WT-TIPs or High-dose CATIPs) or 3 (High-dose CATIPs or positive-control proteins). n.s. *P* > 0.05; **P* < 0.05; *****P* < 0.0001; one-way ANOVA followed by Tukey's multiple comparisons test.

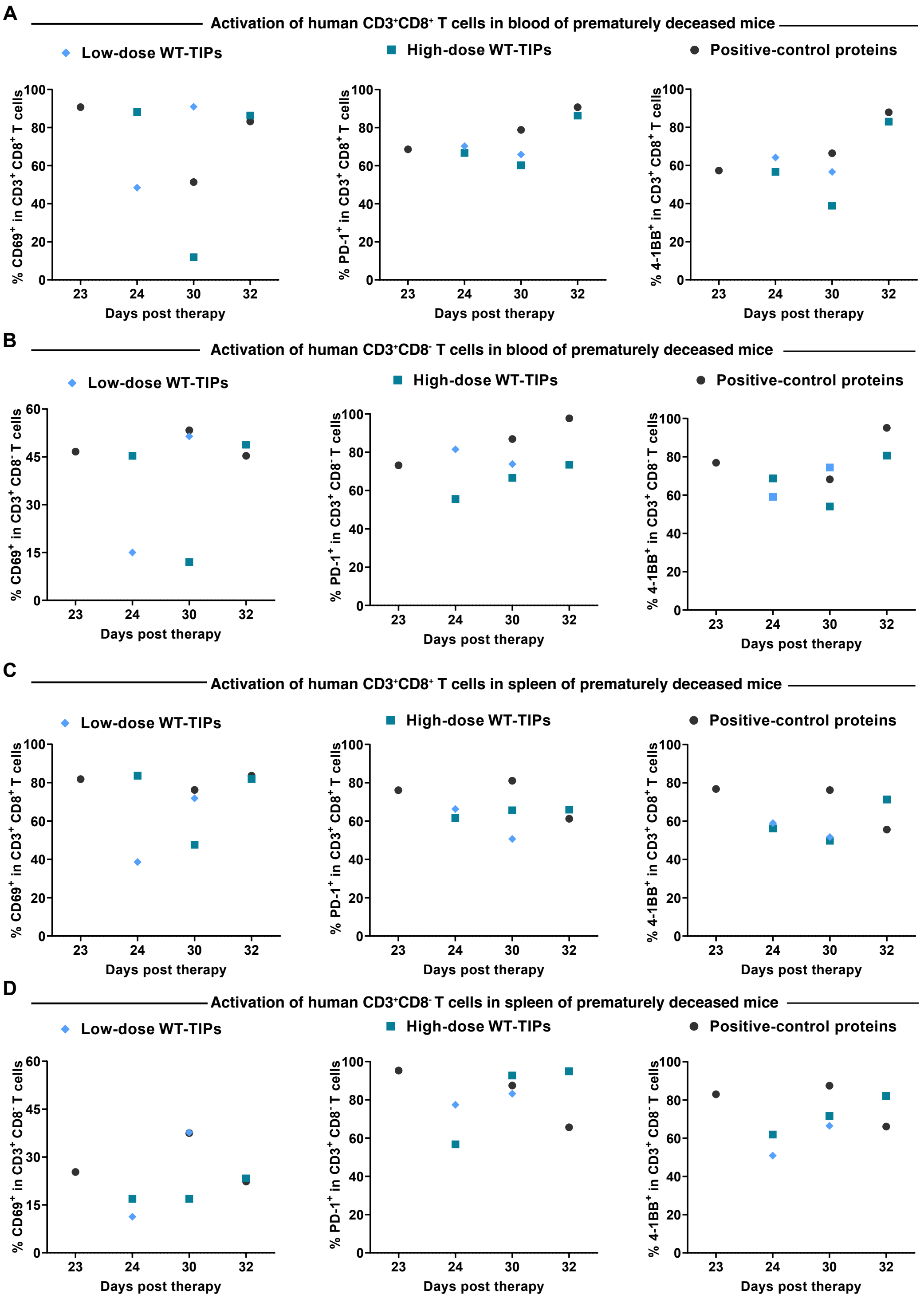


**Figure S5. WT-TIPs induced robust human T cell activation in deceased mice.**

(**A-B**) Activation of human CD3⁺CD8⁺ (A) or CD3⁺CD8⁻ (B) T cells in the blood of deceased mice at the indicated time points.

(**C-D**) Activation of human CD3⁺CD8⁺ (C) or CD3⁺CD8⁻ (D) T cells in the spleens of deceased mice at the indicated time points.

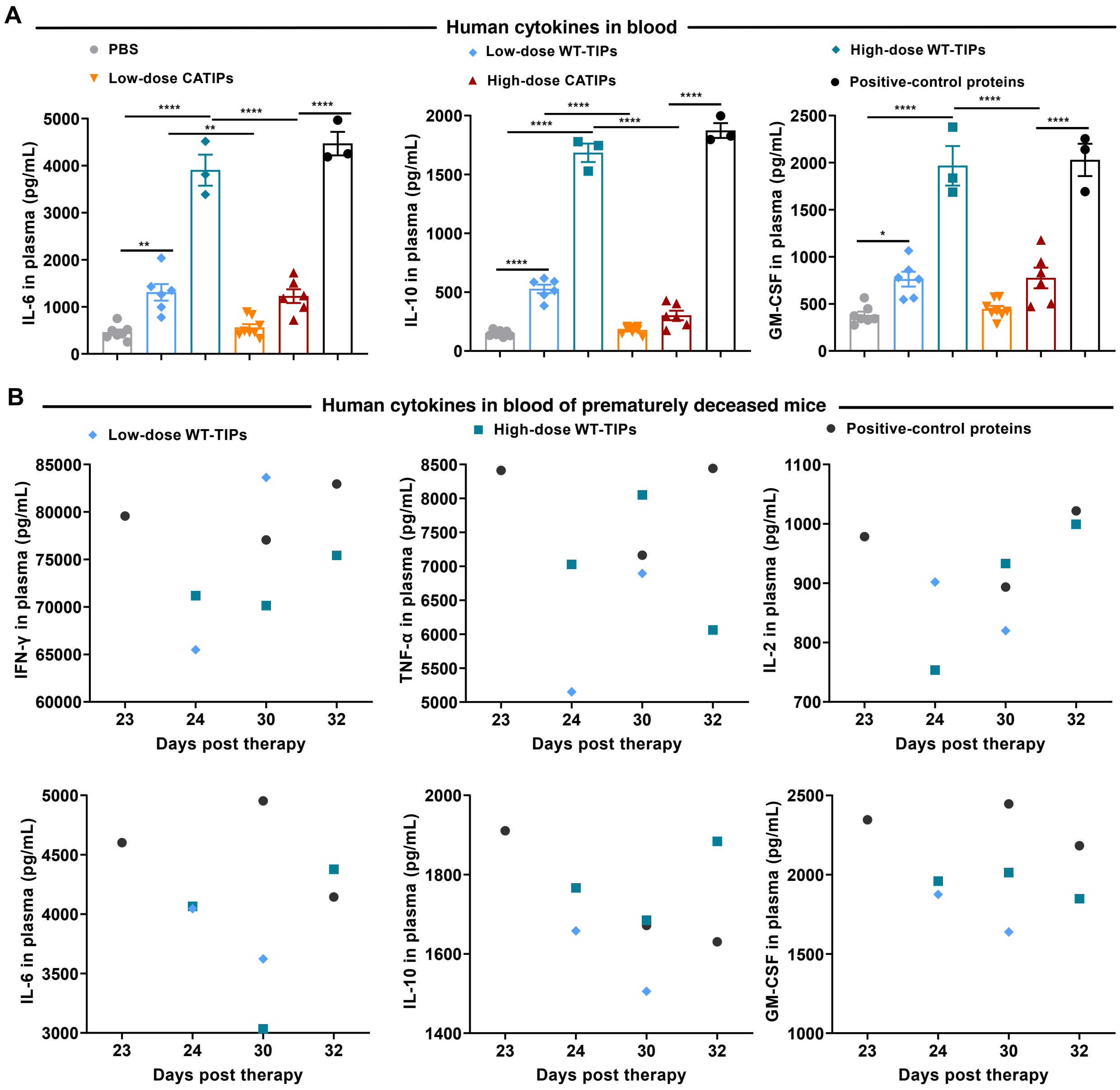


**Figure S6. WT-TIPs induce elevated human cytokine production in deceased mice.**

(**A**) WT-TIPs induced elevated levels of human IL-6, IL-10, and GM-CSF in mouse blood at the study endpoint. Data are mean ± SEM. n = 7 (PBS), 8 (low-dose CATIPs), 6 (low-dose WT-TIPs or high-dose CATIPs), or 3 (high-dose CATIPs or positive-control proteins). **P* < 0.05; ***P* < 0.01; *****P* < 0.0001; one-way ANOVA followed by Tukey's multiple comparisons test.

(**B**) WT-TIPs induced extensive human cytokine levels in deceased mice at the indicated time points.

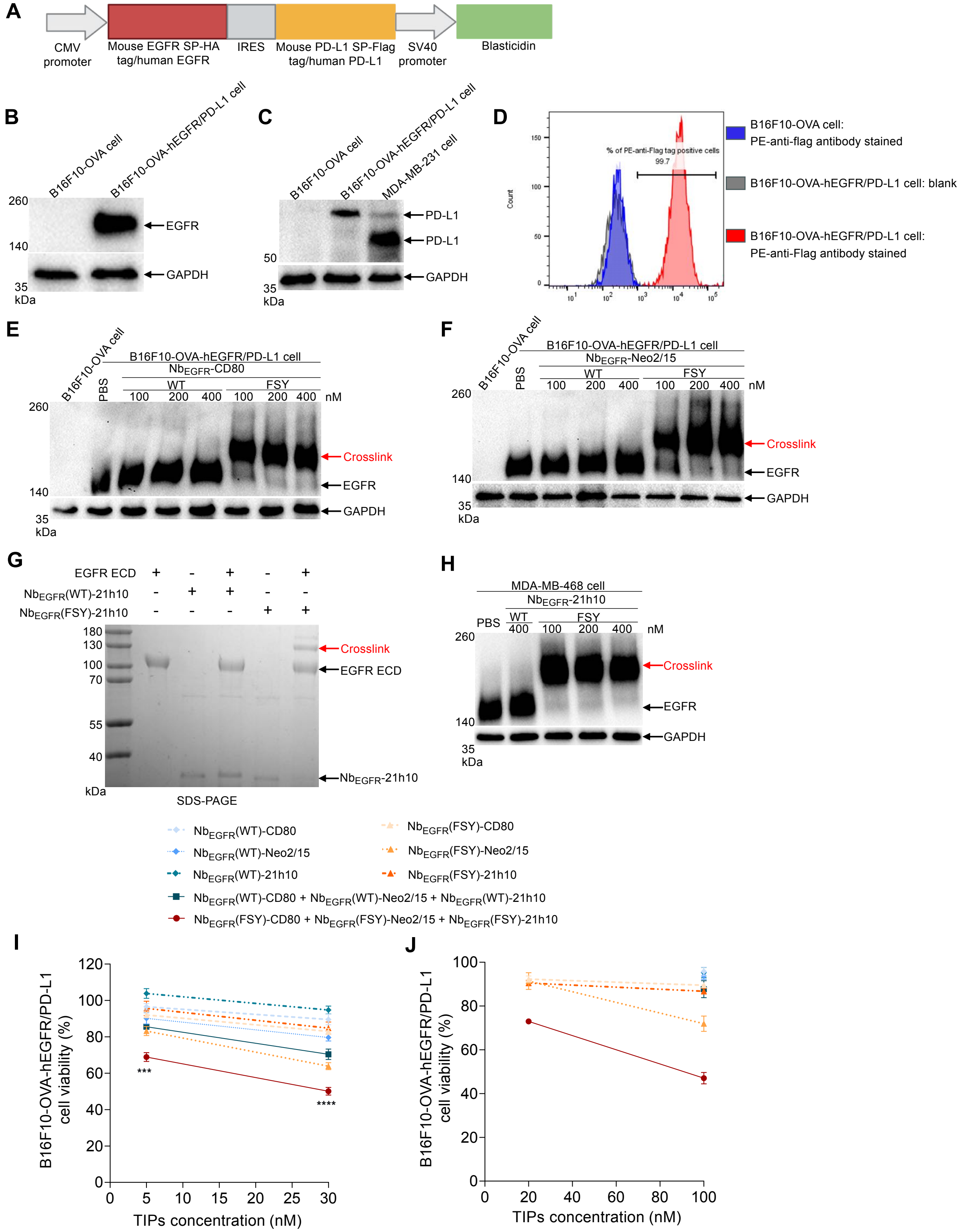


**Figure S7. CATIPs covalently bound EGFR on B16F10-OVA-hEGFR/PD-L1 cells, resulting in enhanced splenocyte-mediated cytotoxicity.**

**(A)** Schematic of the expression construct used to generate B16F10-OVA-hEGFR/PD-L1 cells stably co-expressing human EGFR and PD-L1.

(**B-C**) Western blot analysis of the generated B16F10-OVA-hEGFR/PD-L1 cells stably expressing human EGFR (B) and PD-L1 (C).

(**D**) Flow cytometry analysis of human PD-L1 expression on the surface of B16F10-OVA-hEGFR/PD-L1 cells.

(**E-F**) Western blot analysis of $Nb_{EGFR}$(FSY)-CD80 (E) and $Nb_{EGFR}$(FSY)-Neo2/15 (F) covalently bound EGFR on the surface of B16F10-OVA-hEGFR/PD-L1 cells.

(**G**) SDS-PAGE analysis of $Nb_{EGFR}$(FSY)-21h10 covalently crosslinked EGFR ECD in vitro. $Nb_{EGFR}$-21h10 was incubated with EGFR ECD at a molar ratio 1:1 in PBS buffer at 37°C for 14 h.

(**H**) Western blot analysis of $Nb_{EGFR}$(FSY)-21h10 covalently bound EGFR on the surface of MDA-MB-468 cells.

(**I-J**) CATIP combinations cooperatively enhanced mouse splenocyte-mediated cytotoxicity against B16F10-OVA-hEGFR/PD-L1 cells under PBS non-washed (I) or washed (J) conditions in a dose-dependent manner. Data are mean ± SEM. n = 4. ***$P < 0.001$, ****$P < 0.0001$; asterisks at the indicated positions denote comparisons between CATIPs and WT-TIPs at equivalent doses (two-way ANOVA with Tukey's multiple-comparisons test).

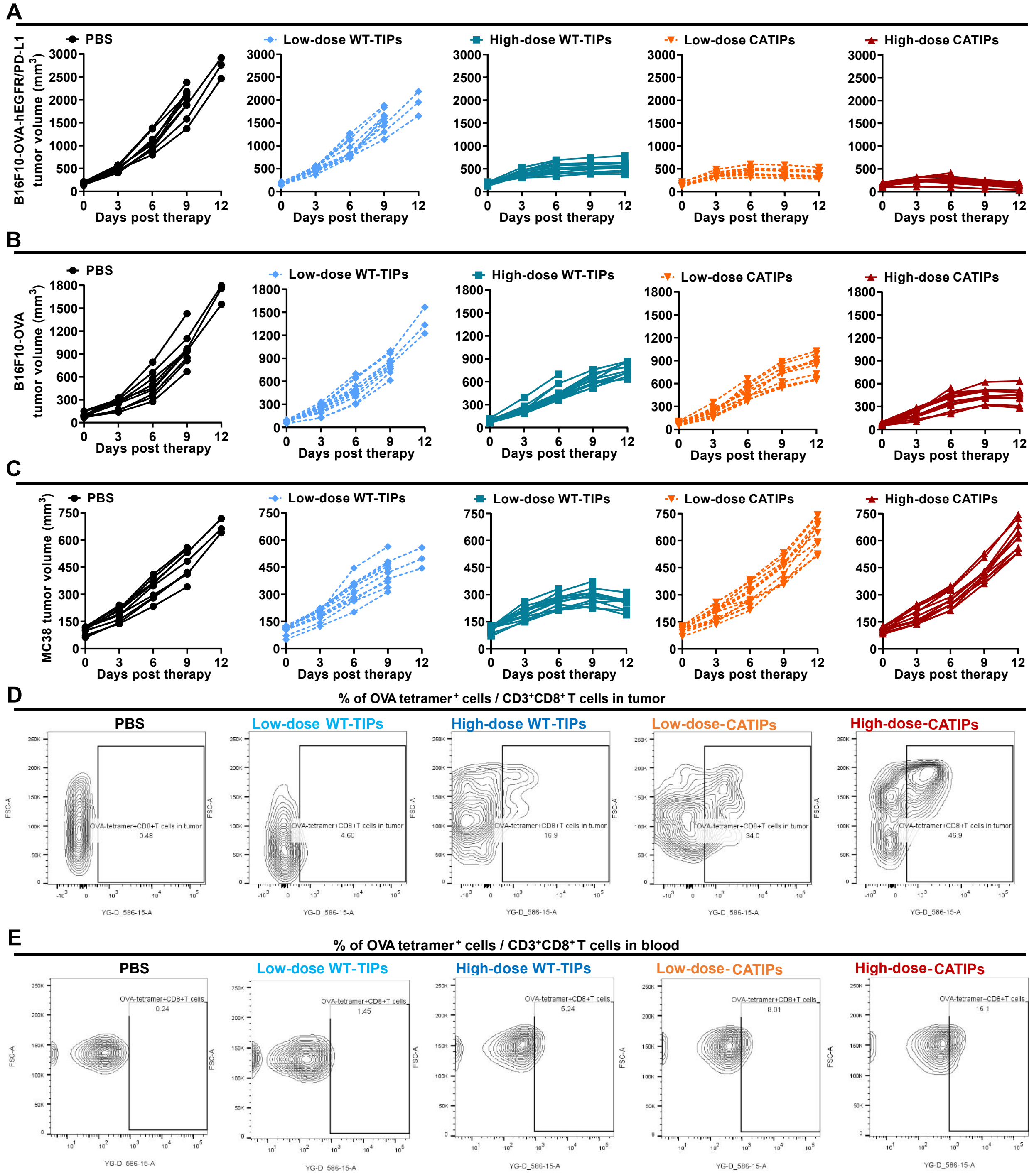


**Figure S8. CATIPs significantly inhibited antigen-specific tumor growth and induced OVA-specific CD8$^{+}$T cells in tumors and blood.**

(**A-C**) Tumor growth curves of B16F10-OVA-hEGFR/PD-L1 tumors (A), B16F10-OVA tumors (B) or MC38 tumors (C) in individual mice.

(**D-E**) Representative flow cytometry plots of OVA tetramer$^{+}$ cells among CD3$^{+}$CD8$^{+}$ T cells in tumors (D) and blood (E).

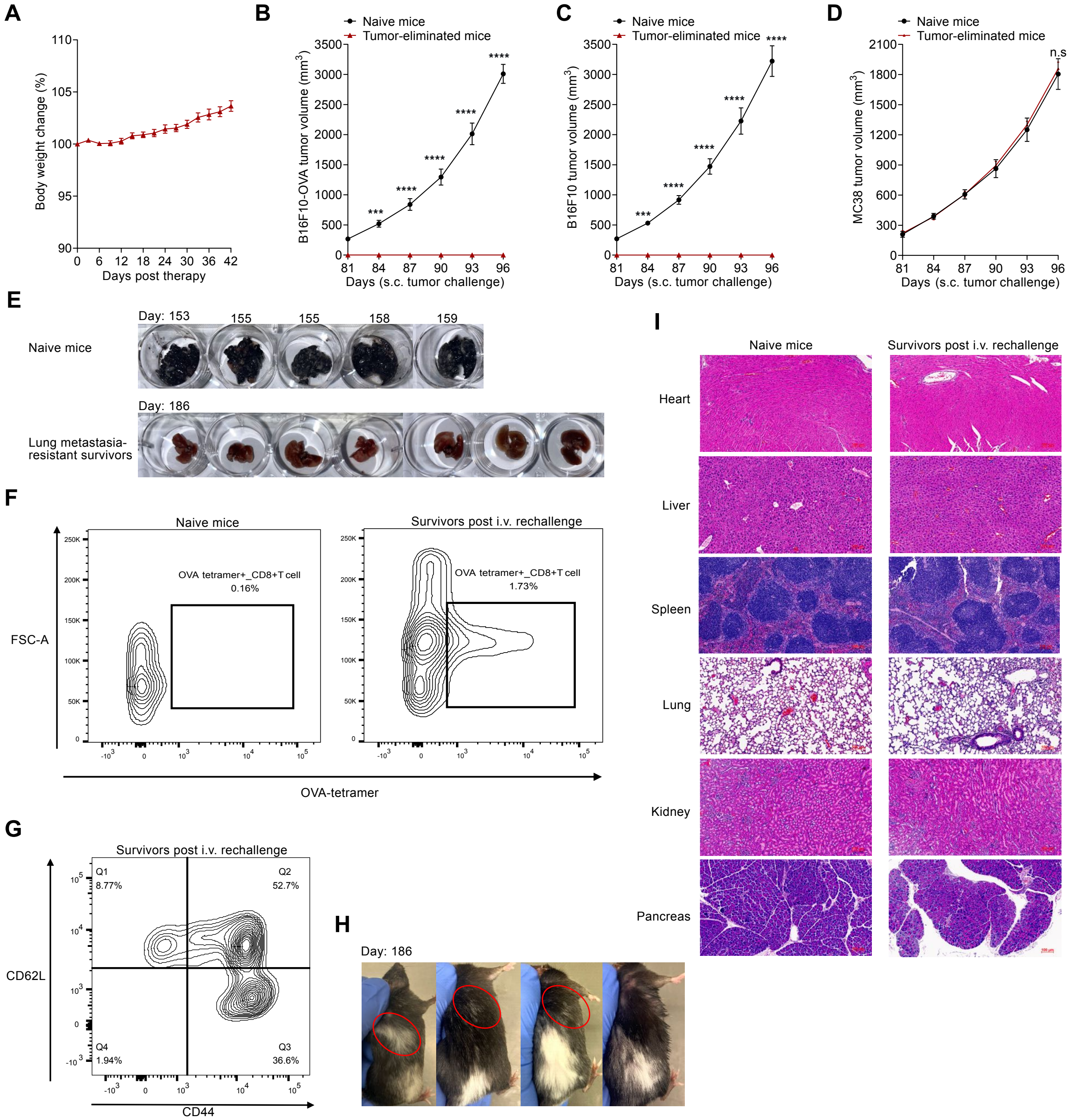


**Figure S9. Antigen-specific immune memory and toxicity in CATIP-treated tumor-eradicated mice.**

**(A)** Relative changes in body weight of mice following high-dose CATIPs treatment.

**(B-D)** Tumor growth curves of B16F10-OVA tumors (B), B16F10 tumors (C) or MC38 tumors (D) in Antigen-specificity model shown in **Figure 5O**. Data are mean ± SEM. n = 4 independent experiments. n.s. $P > 0.05$, ***$P < 0.001$, ****$P < 0.0001$; two-way ANOVA followed by Šídák's multiple-comparisons test.

**(E)** Images of lung metastases following intravenous rechallenge with B16F10-OVA cells at the indicated time points.

**(F)** Representative flow cytometry plots of OVA tetramer$^+$ cells among CD3$^+$CD8$^+$ T cells in harvested spleens at day 186.

(**G**) Representative flow cytometry plots of CD44 CD62L cells among OVA tetramer$^{+}$CD8$^{+}$ T cells in harvested spleens at day 186.

(**H**) Representative images of mice showing vitiligo-like depigmentation with localized hair whitening at the subcutaneous rechallenge site (red circle) observed in 3 of 7 mice. Depigmentation remained stable and confined to the initial treatment site at right posterior flank in all survivors.

(**I**) H&E staining of tissues for evaluation of CATIP-associated toxicity.

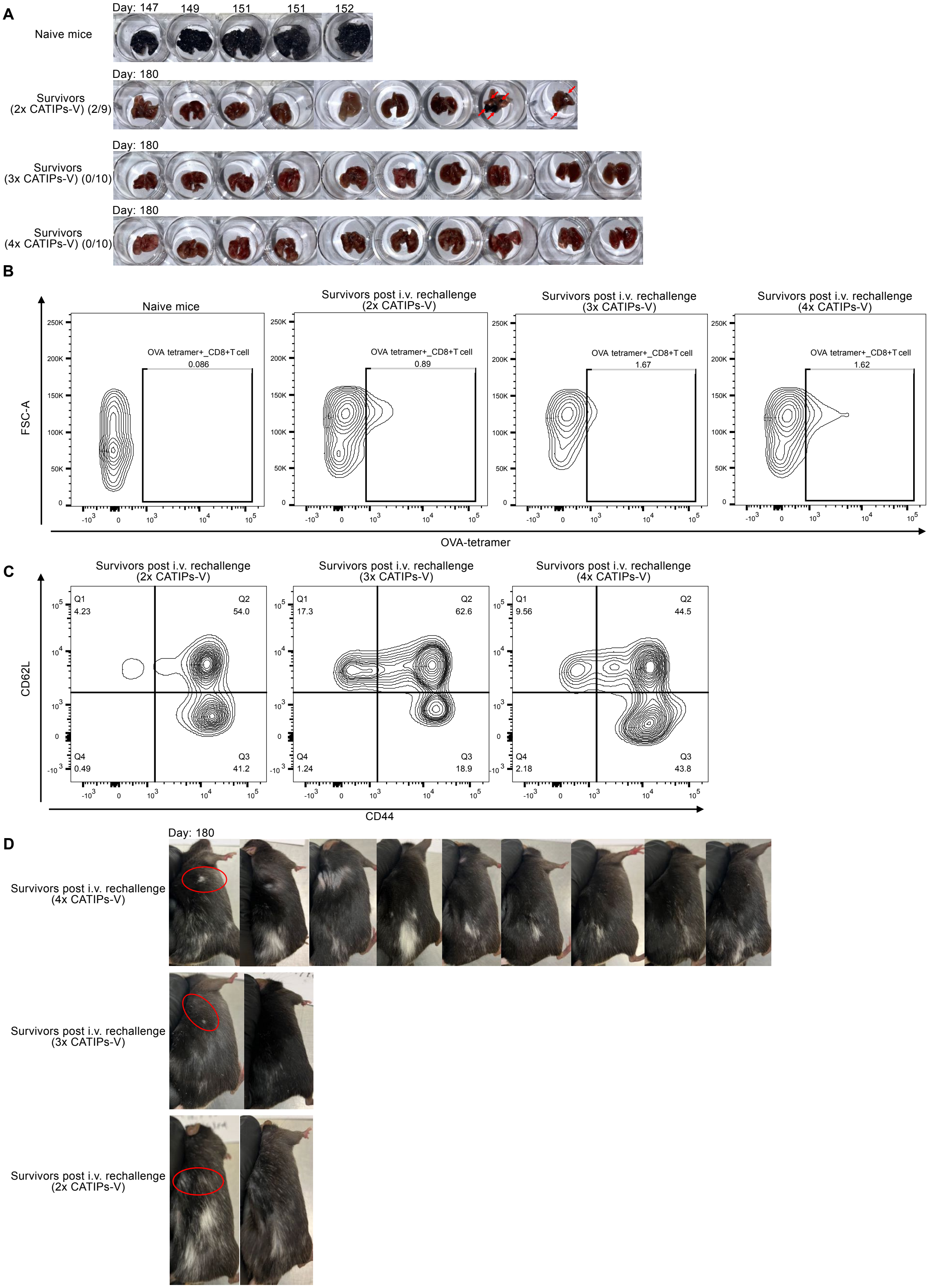
A
Day: 147
149
151
151
152
Naive mice
Day: 180
Survivors
(2x CATIPs-V) (2/9)
Day: 180
Survivors
(3x CATIPs-V) (0/10)
Day: 180
Survivors
(4x CATIPs-V) (0/10)
B
Naive mice
Survivors post i.v. rechallenge
(2x CATIPs-V)
Survivors post i.v. rechallenge
(3x CATIPs-V)
Survivors post i.v. rechallenge
(4x CATIPs-V)
OVA tetramer+_CD8+T cell
0.086
0.89
1.67
1.62
FSC-A
OVA-tetramer
C
Survivors post i.v. rechallenge
(2x CATIPs-V)
Q1 4.23
Q2 54.0
Q4 0.49
Q3 41.2
Survivors post i.v. rechallenge
(3x CATIPs-V)
Q1 17.3
Q2 62.6
Q4 1.24
Q3 18.9
Survivors post i.v. rechallenge
(4x CATIPs-V)
Q1 9.56
Q2 44.5
Q4 2.18
Q3 43.8
CD62L
CD44
D
Day: 180
Survivors post i.v. rechallenge
(4x CATIPs-V)
Survivors post i.v. rechallenge
(3x CATIPs-V)
Survivors post i.v. rechallenge
(2x CATIPs-V)

**Figure S10. Representative data from mice immunized with CATIPs-engineered whole-tumor cell vaccines.**

**(A)** Images of lung metastases following intravenous rechallenge with B16F10-OVA cells at the indicated time points. Red arrows indicate lung metastatic nodules.

**(B)** Representative flow cytometry plots of OVA tetramer$^+$ cells among CD3$^+$CD8$^+$ T cells in harvested spleens at day 180.

**(C)** Representative flow cytometry plots of CD44 CD62L cells among OVA tetramer$^+$CD8$^+$ T cells in harvested spleens at day 180.

**(D)** Representative images of mice showing vitiligo-like depigmentation with localized hair whitening at the subcutaneous rechallenge site (red circle), observed in 1 of 10 survivors immunized with 4x or 3x CATIPs and in 1 of 9 survivors immunized with 2x CATIPs. Depigmentation remained stable and confined to the initial immunization site at right posterior flank in all survivors.
